# From Northwest Passage shores to molecular pathways: Comparative transcriptomic responses of a novel Arctic marine fuel-degrading *Flavobacterium* species

**DOI:** 10.64898/2025.12.09.693246

**Authors:** Nastasia J. Freyria, Antoine-Olivier Lirette, Charles W. Greer, Lyle G. Whyte

## Abstract

Accelerated sea-ice decline is opening the Arctic to increased shipping, elevating the risk of marine fuel spills in fragile ecosystems where extreme cold and remoteness limit cleanup options. While microbial biodegradation is the primary removal mechanism, the metabolic strategies of abundant polar taxa remain poorly understood, particularly those lacking canonical degradation genes. We characterized the hydrocarbon degradation mechanisms of *Flavobacterium* sp. strain R2B_3I, a psychrotolerant isolate from high Arctic beach sediments in Resolute Bay, Nunavut, Canada. During three-month incubations with ultra-low sulfur fuel oil at 4 °C, R2B_3I mounted a systems-level response involving the upregulation of diverse non-canonical oxidoreductases, membrane remodeling systems, cold-shock, and oxidative stress defenses. Crucially, this strain achieved efficient degradation in the complete absence of *alkB* alkane hydroxylases, challenging the reliance on *alkB* as a universal biomarker for hydrocarbon biodegradation. Transcriptomic analysis revealed distinct temporal shifts, linking specific gene clusters to the degradation of complex petroleum mixtures under environmentally relevant conditions. These results demonstrate that *Flavobacterium,* a dominant genus in polar oceans, utilize a “cryptic” metabolic network to process hydrocarbons, effectively bypassing the pathways typically monitored in environmental surveys. By uncovering alternative mechanisms, our study revises current models of microbial oil degradation, highlighting the overlooked potential of non-canonical degraders in determining the fate of marine fuel spills in a warming Arctic.

## INTRODUCTION

The Arctic Ocean is undergoing rapid transformation as climate change is accelerating sea-ice retreat and driving a shift toward thinner, more mobile first-year ice, expanding seasonal access to remote Arctic waters for navigation [1–3]. Arctic sea-ice extent has declined by about 13% per decade since 1979 [1], and seasonally ice-free summers are projected to occur regularly by mid-century [4, 5]. Projections suggest that the Northwest Passage will become increasingly navigable, and that up to 5% of global shipping may be diverted to the Northern Sea Route by 2050 [6], accompanied by rising traffic along shallow shelf seas such as the Beaufort and Chukchi [7]. This expansion in maritime activity increases spill risk in environments where extreme cold, darkness, remoteness, and harsh weather reduce fuel fluidity and limit the effectiveness of mechanical recovery, dispersants, and in-situ burning, leaving natural attenuation as a critical process when engineered interventions are constrained [8–10].

Arctic environments already receive hydrocarbons from both natural and anthropogenic sources [11]. Natural oil seeps have been reported in several Arctic regions [12, 13], including the Beaufort Sea, where seepage releases petrogenic hydrocarbons (e.g., alkanes and aromatics) to nearby sediments and shorelines [14]. Additional inputs arise from biogenic hydrocarbon production by microorganisms [15], sea-air exchange along coastal zones [16], and atmospheric deposition [17]. In contrast, the growing risk of marine fuel oil releases from shipping represents a distinct contamination pathway that warrants investigation of degradation under polar conditions. Ultra-low sulfur fuel oil (ULSFO), in particular, is a complex mixture of hydrocarbons from roughly C10 to C50, with a high proportion of aromatics and resin fractions that are especially recalcitrant in cold environments and therefore more likely to cause persistent, long-term impacts [18, 19].

Arctic microbiomes host cold-adapted microbial communities active under extreme conditions, including seawater temperatures as low as −4 °C to −2 °C, atmospheric temperatures ranging from −40 °C to 10 °C, with up to six months of darkness and strong physicochemical variability. Psychrophiles (optimal growth below 15 °C, typically 10-15°C, with growth range from −15 °C to 20 °C, no growth above 20 °C) and psychrotolerant taxa (capable of growth at 0 °C, optimal growth at 20-30 °C, with growth range from −5°C to 35-40 °C) exhibit adaptations such as altered membrane lipid composition for enhanced fluidity [20, 21], cold-shock and RNA chaperone systems [22, 23], and modified enzyme kinetics that sustain activity at low temperatures, despite slower overall metabolic rates relative to mesophiles [24, 25]. Multiple bacterial genera play established roles in hydrocarbon degradation at low temperature, including *Pseudomonas* (n-alkanes) [26, 27], *Rhodococcus* (aromatic compounds) [28–31], and *Alcanivorax* (alkane specialists) [32, 33]. *Flavobacterium* is broadly distributed in polar environments, including Arctic soil [34, 35], glaciers [36, 37], freshwater sediments [38], Antarctic lakes [39], sea-ice [40], Arctic seawater and coastal sediment environments [41–43]. Although many *Flavobacterium* strains have been isolated from cold environments, relatively few studies have demonstrated their ability to metabolize diverse hydrocarbons, including n-alkanes and polycyclic aromatic hydrocarbons (PAHs), in-situ or in culture, with examples from oil-contaminated Arctic soils [34, 44, 45] and Subantarctic coastal sediments [46]. Genomic and transcriptomic evidence for key alkane hydroxylases (*alkB*, *almA*, *ladA*), PAH-degrading dioxygenases, and central aromatic pathways (*catABC*, *pca*) remains inconsistent [28, 47, 48], while anaerobic activation genes (*assA/masD*, *bssA*) are rarely detected despite oxygen limitation in Arctic sediments [49]. To date, most investigations have employed simple model substrates or crude oils, with far less attention to the heavy marine fuels that now dominate Arctic shipping. Consequently, how *Flavobacterium* processes ULSFO under realistic polar conditions remains largely unresolved, limiting our ability to predict natural attenuation or design targeted bioremediation strategies.

Here, we investigate the biodegradation potential of indigenous *Flavobacterium* from Arctic beach sediments under controlled laboratory conditions simulating Arctic marine environments (4 °C, continuous darkness) for three months with ULSFO exposure. We paired cultivation with comparative transcriptomics to elucidate molecular mechanisms underlying hydrocarbon processing, focusing on upregulated genes encoding hydrocarbon-degrading enzymes, stress responses, and cold-adaptive metabolism that enable growth on complex petroleum mixtures at low temperature. We hypothesize that ULSFO exposure induces specialized, cold-adapted hydrocarbon pathways in *Flavobacterium* involving novel combinations of alkane activation, aromatic degradation, and cold-shock systems not previously characterized in psychrophilic degraders. These insights aim to inform enhanced bioremediation strategies and improve forecasts of natural attenuation capacity in Arctic marine ecosystems facing increased spill risk from expanding shipping.

## MATERIALS & METHODS

### Strain isolation and growth conditions

*Flavobacterium* sp. strain R2B_3I was isolated from Arctic beach sediment (Tupirvik Beach, Resolute Bay, Nunavut, Canada) following established protocols [29]. Briefly, 5 g of sediment from a mesocosm simulating intertidal cycles [27] were suspended in sterile seawater (1:4 w/v) with glass beads, vortexed to dislodge particles, and serially diluted. Aliquots were spread onto R2A gellan plates and minimal media plates amended with 500 ppm ultra-low sulfur fuel oil (ULSFO) as the sole carbon source.

Hydrocarbon biodegradation experiments were conducted in 600 mL sterile artificial seawater (3% sea salt) supplemented with 0.67 g.L□^1^ ammonium sulfate. Treatment flasks received 500 ppm ULSFO (~45 mg per 100 mL). Triplicate flasks were incubated under four conditions: 1) abiotic control (ULSFO without cells); 2) *Flavobacterium* biotic control (cells in R2A broth without ULSFO); and 3) oil-exposed cells harvested at 1-month (T1) and 4) 3-months (T3). All cultures were inoculated at an initial density of 1×10 cells.mL□^1^ and shaken at 100 rpm at 4 °C in the dark. At T0, T1, and T3, cells were harvested by centrifugation (5,000 rpm, 2 min), resuspended in DNA/RNA Shield (Zymo Research), and stored at −80 °C.

### Petroleum hydrocarbon analyses

At each sampling point, culture media were warmed to 60 °C to resuspend ULSFO adhering to flask walls before transfer to glass bottles and storage at −20 °C. Semi-volatile organic compounds (SVOCs) were extracted via liquid-liquid extraction with dichloromethane and quantified using US EPA Method 8270E [50]. Petroleum hydrocarbon, including polycyclic aromatic hydrocarbons (PAHs) and aliphatic fractions (F1-F4), were analyzed following CCME Reference Method protocols [51]. Initial concentrations were established from abiotic controls. Hydrocarbon removal was calculated as the difference between initial and residual concentrations (sum of oil- and water-phase), expressed as a percentage of the initial ULSFO concentration. All conditions were analyzed in biological triplicate with technical duplicates (n = 6 per condition).

### Whole genome sequencing and annotation

Genomic DNA was extracted using DNeasy PowerSoil kit (Qiagen). DNA quantity and purity were assessed using a Qubit fluorometer (dsDNA kit) and NanoDrop 8000 spectrophotometer. Sequencing was performed on an Illumina NovaSeq 6000 S4 platform (paired-end 150 bp) at Genome Québec. The assembled genome (5.16 Mb; 6,028 genes) was processed by trimming reads with Trimmomatic v0.36 [52] and assembled using SPAdes v3.15.4 [53]. Functional annotation was performed with METAerg [54], and hydrocarbon degradation genes were identified using CANT-HYD [55] (E-value <0.01). The assembled genome was submitted to the JGI Integrated Microbial Genomes database [56].

Phylogenetic trees were constructed from curated full-length 16S rRNA sequences using RAxML v8.2.11 [57] (GTRGAMMA model, 1,000 bootstraps) following MUSCLE alignment. To resolve evolutionary relationships of degradation genes, gene trees were constructed for *alkB*, *almA*, and *ompW* using appropriate substitution models. Putative homologs were identified by blastp against specific reference proteins: *alkB* against *Flavobacterium* sp. 1-C-1 (ACJ22756.1); *almA* against *Alloalcanivorax dieselolei* (ADP30851.1); and *ompW* against *Flavobacterium sinopsychrotolerans* (CAM3936058.1). Pangenome analysis of 11 *Flavobacterium* genomes was performed with Roary v3.13.0 [58] with Prokka annotation [59]. Average nucleotide identity was computed with fastANI v1.34 [60].

### Transcriptomic sequencing and processing

Total RNA was extracted using RNeasy Mini Kit (Qiagen) with quality assessment via Qubit fluorometer and NanoDrop spectrophotometer. Libraries were prepared using the NEBNext rRNA Depletion Kit (Bacteria) and sequenced on Illumina NextSeq platform (paired-end 100 bp), generating ~583 million reads. We analyzed 15 samples: three biological replicates each at T0, T1, T3 for biotic controls, and T1 and T3 for ULSFO-exposed cells. Raw reads were trimmed with BBMap v38.18 [61] and mapped to the R2B_3I reference genome using STAR v2.7.10b [62]. Gene-level counts were quantified across biological replicates and pooled (**Table S1**). Functional annotations (GO, KEGG, COG) were retrieved from JGI Genome Portal. CAZymes were predicted with dbCAN [63] and hydrocarbon markers with CANT-HYD.

### Co-expression networks

Weighted Gene Co-expression Network Analysis (WGCNA) [64] was used to assess ULSFO exposure effects over time, as previously described [65, 66]. Networks were constructed with soft-thresholding power of 15, minimum module size of 50 genes, and merge cut height of 0.25. Module-trait relationships were determined through Pearson correlations between module eigengenes and experimental conditions. Hub genes were identified based on high intramodular connectivity and module membership.

### Statistical analyses

Statistical analyses were conducted in R v4.3.3. Gene counts were normalized using DESeq2 [67] (median ratio method). Differentially expressed genes (DEGs) were defined as those with adjusted p ≤0.05 and |log FC|≥2. DEG representations were visualized with scatter plots as described previously [68]. Two-way repeated-measures ANOVA using PAST v4.11 to assess time-by-condition effects for petroleum hydrocarbons. Canonical correspondence analysis was performed using vegan::cca [69]. Heatmaps were generated from variance-stabilized counts using pheatmap [70] and circlize [71].

### Data availability

Transcriptome sequencing datasets are available at NCBI (BioProject: PRJNA1193011). Whole genome sequencing is available on JGI portal (Gold Analysis Project ID: Ga0555976) and NCBI (BioProject: PRJNA945214).

## RESULTS

### Evolutionary diversification reveals *Flavobacterium* sp. strain R2B_3I as a novel Arctic lineage

Phylogenetic analysis of 113 *Flavobacterium* strains identified five distinct clades (I-V), each supported by bootstrap values >50 (**Fig. 1A**). Strain R2B_3I grouped within Clade V alongside *F. gillisiae*, *F. frigidarium*, and *F. psychrophilum*. However, low 16S rRNA sequence identity (<97%) and comparative genomic analysis clearly indicated R2B_3I represents a distinct genomic lineage. Pangenome analysis of 12 Clade V genomes revealed pronounced genomic plasticity: unique genes comprised 75.8% (14,802 genes) of the total, while core genes represented only 2.9% (573 genes), with shell genes (11%), cloud genes (7.9%), and soft-core genes (2.3%) indicating variable genome content (**Fig. 1B**). Average nucleotide identity (ANI) analysis showed highest similarity with *F. degerlachei* (81.44%) and *F. frigoris* (81.22%), well below species boundaries (**Fig. 1B, Table S2**). Even the most similar pair among known genomes (*F. frigoris* and *F. degerlachei)* exhibited an ANI of only 88.40%, further underscoring the deep evolutionary divergence within this clade. Collectively, the low genomic relatedness, phylogenetic distinctness, and abundance of unique genes strongly support that R2B_3I represents a novel species within *Flavobacterium*.

**Fig. 1.**
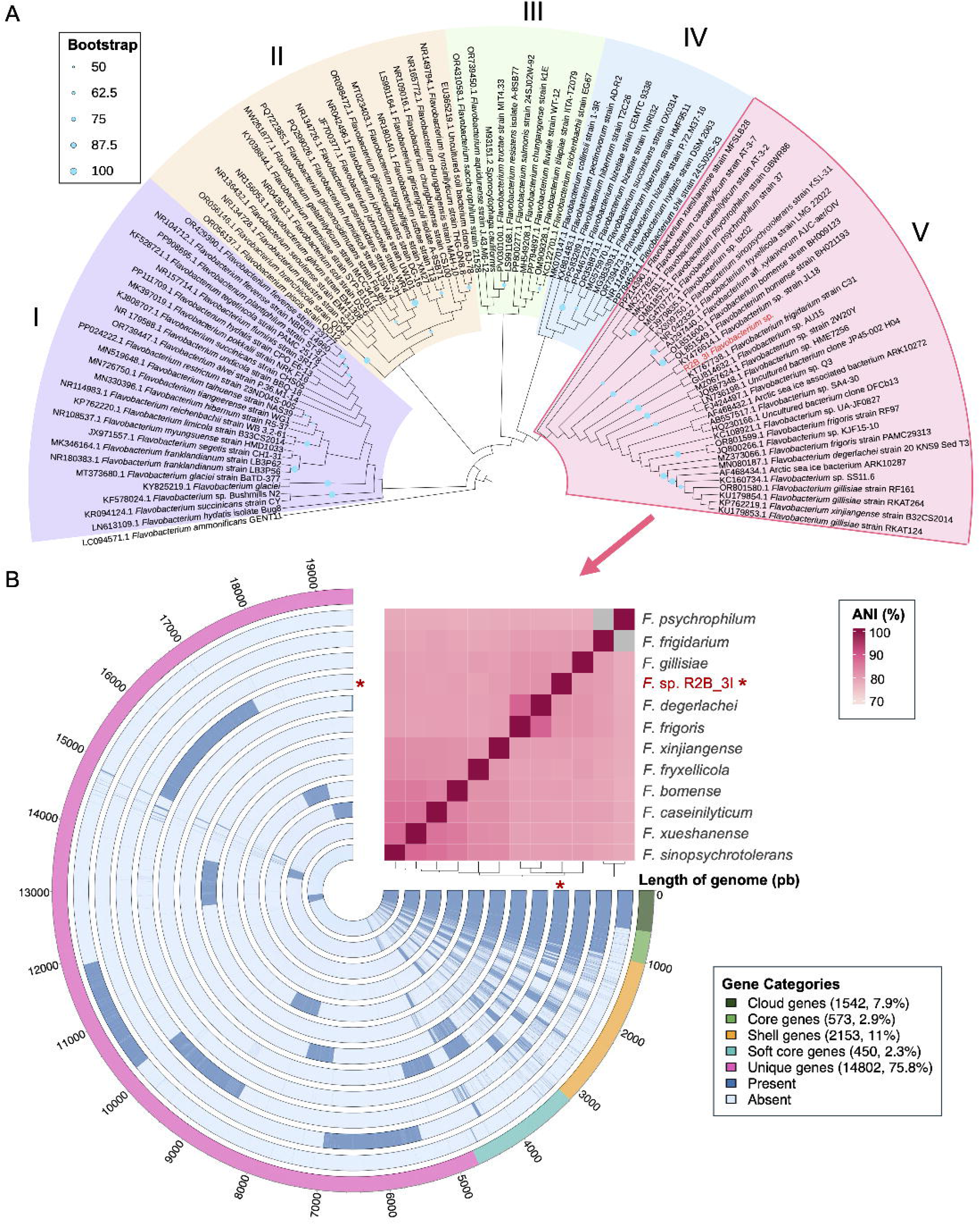
A Maximum Likelihood *Flavobacterium* tree and a pangenome of 12 *Flavobacterium* genomes. **(A)** A total of 112 *Flavobacterium* strains and species 16S rRNA gene sequences were aligned along with one reference sequence of a short length 16S rRNA gene (sequenced using the Sanger method). The phylogenetic tree was constructed using RAxML from an alignment of 113 sequences, spanning 1400 characters, with bootstrap support calculated over 1,000 repetitions. Only bootstrap values greater than 50 (out of 100) are displayed. Numbers on top of the tree (I to V) refer to different clade. (**B)** The circular pangenome compares 12 *Flavobacterium* genomes from clade V, and the heatmap on top of the pangenome refer to the average nucleotide identity (ANI) between each genome. Core gene: genes present in all genomes; cloud gene: genes present in less than 15% of genomes; shell gene: genes present in 15-95% of genomes; and unique gene: genes present in only 1 genome.

### Petroleum hydrocarbon analysis during ULSFO exposure

Over the three-month exposure, both alkane and aromatic fractions showed distinct temporal patterns (**Figs. S1-S2; Table S3**). Alkanes were resolved into four chain-length fractions, with medium-chain alkanes (F3) dominating at 78,000 μg.L□^1^ at T0 and declining slightly thereafter, while short-chain (F2) and long-chain (F4) alkanes remained comparatively low (>30, 000 μg.L□^1^; **Fig. S1**). The aromatic fraction comprised 19 PAHs, with pyrene, chrysene, and benzo[a]anthracene at the highest concentrations (200-600 μg.L□^1^). Removal efficiencies varied among compounds comparing a single replicate (T3 R1; **Fig. S2, Table S3**): low-molecular weight PAHs, including naphthalene (91.88% removal), fluorene (25.71%), acenaphthene (26.36%), and methylnaphthalenes (63-70%) showed positive removal, while higher-molecular-weight PAHs exhibited apparent increases, likely from desorption or transformation processes. Statistical analyses using one-way ANOVA across the experimental timeline (T0: start, T1: 1 month, T3: 3 months) identified significant temporal changes only for the short-chain alkanes F1 (C6-C10; F = 407.2, *p* <0.001), with t-test and Wilcoxon test confirming significant changes over time for both total aromatic and aliphatic compounds (*p* <0.001 for each; **Table S4**).

### Differential gene expression reveals phased transcriptional response to oil

Differential gene expression analysis from 15 transcriptomes (in triplicate conditions) revealed clear temporal and treatment-specific patterns (**Fig. 2**). Principal component analysis (PCA) showed strong separation of samples by both treatment and time, with PC1 (45.4% variance) primarily separating oil versus control conditions and PC2 (16%) capturing temporal variation (**Fig. 2A**). Hierarchical clustering analysis corroborated the PCA results, with two major clusters separating oil versus control conditions, and sub-clustering by time points (**Fig. 2B**). The heatmap of the 100 most differentially expressed genes (DEGs) further highlighted treatment-specific transcriptional signatures, with consistent biological triplicate grouping (**Fig. 2C**). The hierarchical clustering of genes (vertical axis) revealed six major functional clusters (a-f, detailed in **Table S5**).

**Fig. 2.**
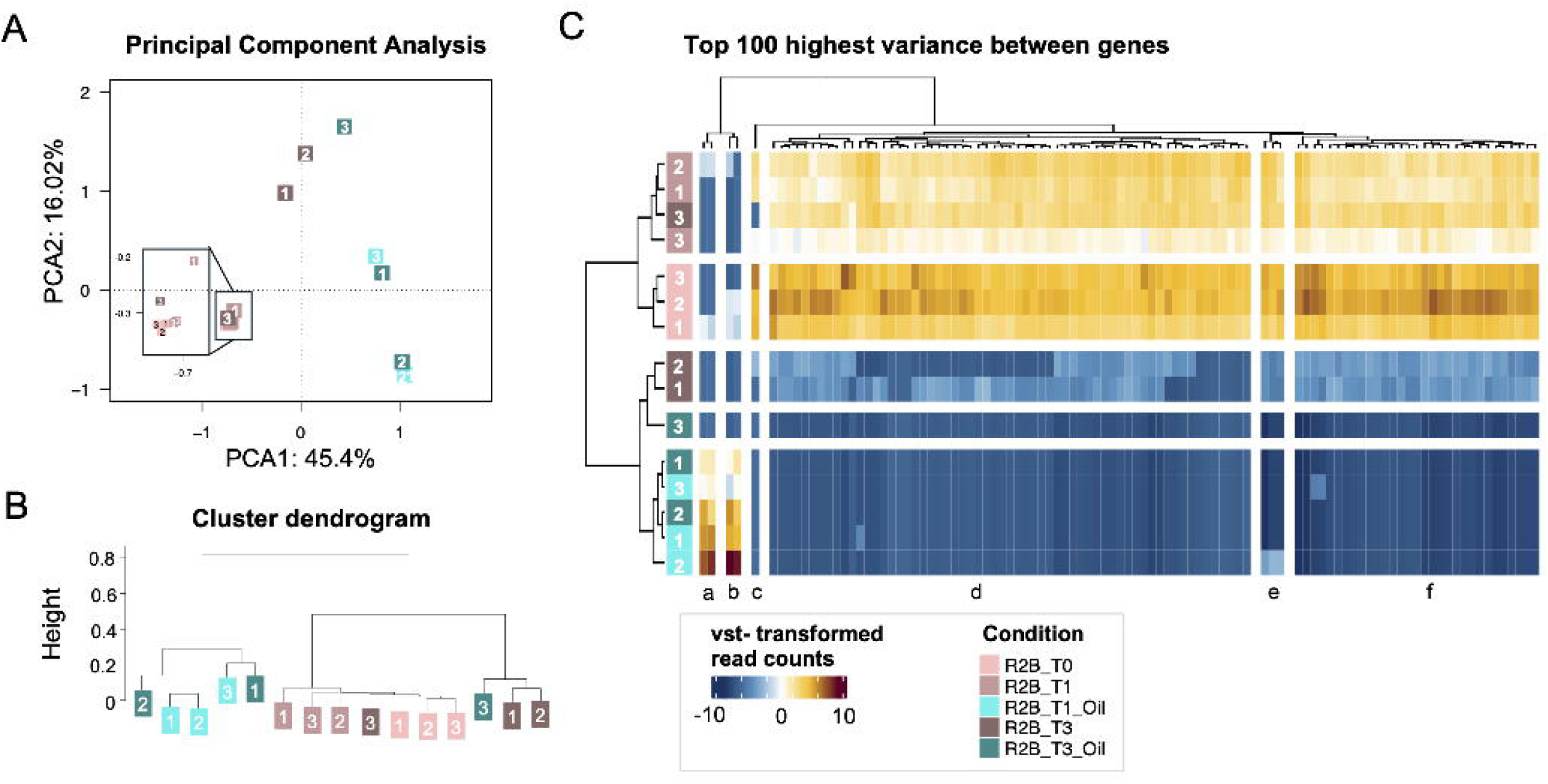
Multi-level analyses of transcriptome responses to oil treatment. **(A)** Principal Component Analysis (PCA) of gene expression profiles using Bray-Curtis dissimilarity measure. Sample conditions are indicated by colors. **(B)** Hierarchical cluster dendrogram of all samples based on normalized read counts, constructed using the “complete” linkage method in the hclust() function (Vegan package, R). Numbers indicate biological replicates. **(C)** Heatmap showing expression patterns of the top 100 differentially expressed genes. Color scale represents variance-stabilized transformed (vst) read counts, calculated using the DESeq2 package. Genes are clustered into functional groups (a-f, detailed in **Table S5**). Dendrograms show hierarchical clustering of both samples (top) and genes (right). Sample labels indicate treatment conditions and biological replicates.

Oil exposure induced a contrast transcriptional change compared to controls (**Fig. S3**). At T1, oil samples showed 3,102 DEGs with 755 significantly upregulated (with adjusted *p-*value <0.05) versus controls with 1,939 DEGs and 1,770 significantly upregulated, reflecting extensive early metabolic reprogramming. By T3, differences were more focused: 972 DEGs with only 168 significantly upregulated in oil versus 1,940 DEGs at T0 with 1,743 significantly upregulated. Direct temporal comparison (T1 oil vs. T3 oil) revealed 2,766 DEGs upregulated at T1 versus only 4 at T3, indicating transition from acute response to sustained adaptation. Among the four T3-upregulated genes were a transposase (cog1943, K07491, pf01797) and L,D-transpeptidase (pf13645), suggesting ongoing genomic flexibility and cell wall remodeling during prolonged exposure.

### Co-expression network analysis reveals distinct response modules

We first explored the expression patterns of 5,069 DEGs detected in all transcriptomes using Weighted Gene Co-expression Network Analysis (WGCNA). The analysis classified these genes into nine co-regulated modules according to the similarity of expression patterns of each gene in the hierarchical clustering dendrogram (**Fig. 3A-B**). Module-trait relationship analysis revealed significant correlations between specific modules and experimental conditions, confirming biological rather than technical clustering (**Fig. 3C**). The Brown module showed strong positive correlation with baseline control (r = 0.87, *p* <0.001), Black and Blue modules correlated with early oil response (r = 0.71, and r = 0.72, *p* <0.01), and Grey module with late oil treatment (r = 0.55, *p* <0.03). Module membership analysis demonstrated strong correlations between gene significance and module membership, particularly in the Brown module (Cor = 0.32, *p* = 1.2e-48) and Blue module (Cor = 0.17, *p* = 1.2e-20; **Fig. 3D**), indicating that condition-relevant genes are most connected within their modules, a hallmark of biologically meaningful co-expression networks.

**Fig. 3.**
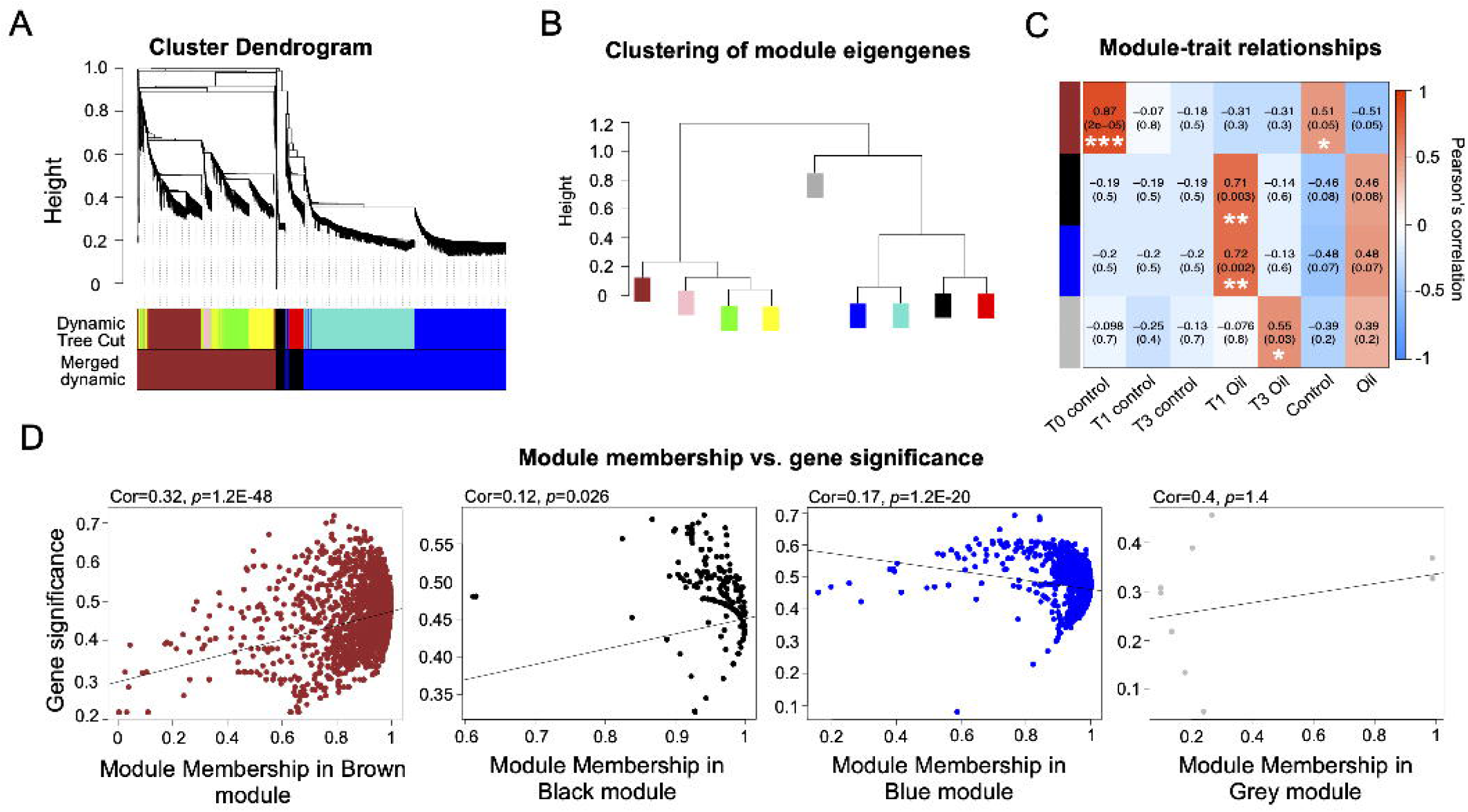
Weighted Gene Co-expression Network Analysis (WGCNA) reveals modular organization of transcriptional response to oil exposure. **(A)** Hierarchical clustering dendrogram of differentially expressed genes (DEGs) showing module assignments (color bands). Clustering based on topological overlap dissimilarity measure (1-TOM). **(B)** Module relationship dendrogram showing hierarchical clustering of module eigengenes, revealing higher-order relationships between response modules. **(C)** Module-trait relationship heatmap showing correlations between module eigengenes and experimental conditions. Color scale represents Pearson correlation coefficients; text shows correlation value and *p*-value in parentheses. Asterisks indicate significance levels: *p <0.05, **p <0.01, ***p <0.001. **(D)** Scatter plots showing correlation between module membership and gene significance for four key modules (brown, black, blue, and grey). Each point represents a gene; trend lines show linear relationship; Cor = correlation coefficient, p = significance value.

### Multi-database functional analysis reveals comprehensive metabolic restructuring

To comprehensively characterize the molecular response to oil exposure, we employed multiple complementary databases that each provide unique insights into different aspects of cellular function. Kyoto Encyclopedia of Genes and Genomes ontology (KEGG) pathway analysis revealed the strongest responses at early exposure (T1; **Fig. 4, Table S6**). In T1 control vs. T1 oil comparison, central carbon metabolism was markedly upregulated in oil samples (glycolysis/gluconeogenesis: 119 DEGs; TCA cycle: 88 DEGs; log2FC <-2), alongside xenobiotic degradation pathways (naphthalene/anthracene: 59 DEGs; benzoate: 48 DEGs). In T3 control vs. T3 oil comparison confirmed elevated expression at T1, while T3 showed reduced xenobiotic degradation (naphthalene/anthracene: 16 DEGs; benzoate: 22 DEGs) but sustained central metabolism (glycolysis/gluconeogenesis: n = 98 DEGs; TCA cycle: n = 74 DEGs; log2FC >2). Energy-related pathways (oxidative phosphorylation, nitrogen/sulfur metabolism) and lipid metabolism showed consistent modulation across timepoints.

**Fig. 4.**
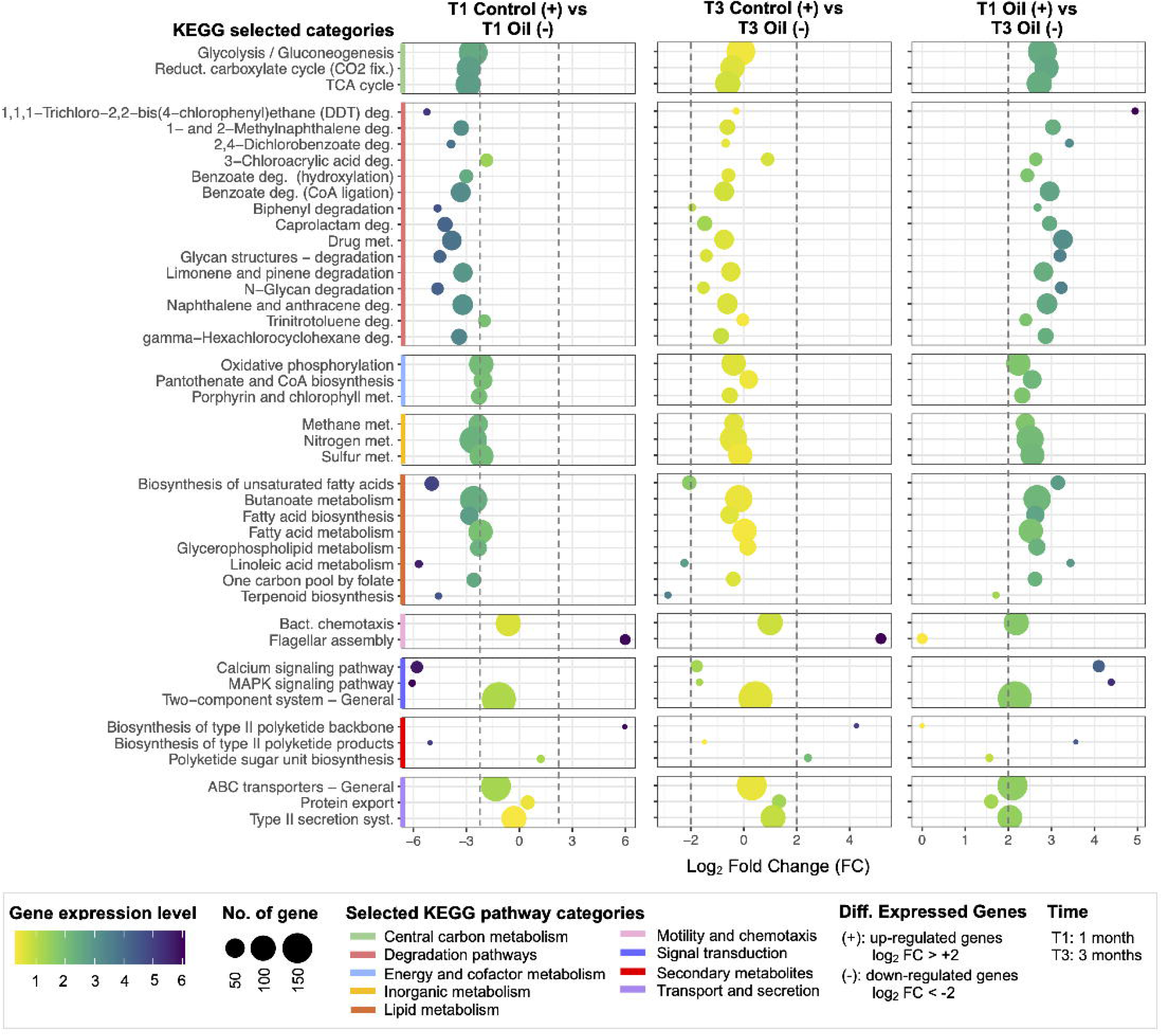
Differential expression analysis of KEGG pathways in response to Ultra-Low Sulfur Fuel Oil (ULSFO) exposure. The dot plot shows log2 fold changes (FC) of differentially expressed genes (DEGs) across three comparisons: T1 Control vs T1 Oil, T3 Control vs T3 Oil, and T1 Oil vs T3 Oil. Dot size indicates the number of genes in each pathway, and color intensity represents gene expression level. Pathways are grouped by functional categories indicated by colored bars. Significant differential expression was defined as log2FC >2 for upregulation and log2FC <-2 for downregulation. Complete gene lists and annotations are available in **Table S6.**

While KEGG provided metabolic pathway context, we next examined functional gene categories using Clusters of Orthologous Groups (COG) to understand broader cellular processes and Carbohydrate-Active enZymes (CAZy) to specifically assess carbohydrate metabolism, both critical for understanding how bacteria modify their cellular machinery and process complex hydrocarbons. COG category analysis revealed distinct temporal patterns in cellular responses to oil (**Figs. 5, S4, Table S7-S8**). At T1 oil, upregulation occurred in carbohydrate/coenzyme transport and metabolism (categories G, H), translation (J), cell wall biogenesis (M), protein modification (O), and defense mechanisms (V; **Fig. 5A**). By T3, most categories showed moderate expression. CAZy family analysis demonstrated extreme upregulation (log2FC <-10) of glycosyltransferase GT84 at T1, with glycoside hydrolases showing varied patterns, GH92 highest at T1 (log2FC >6) and GH37 upregulated only at T3 (**Fig. 5B**).

**Fig. 5.**
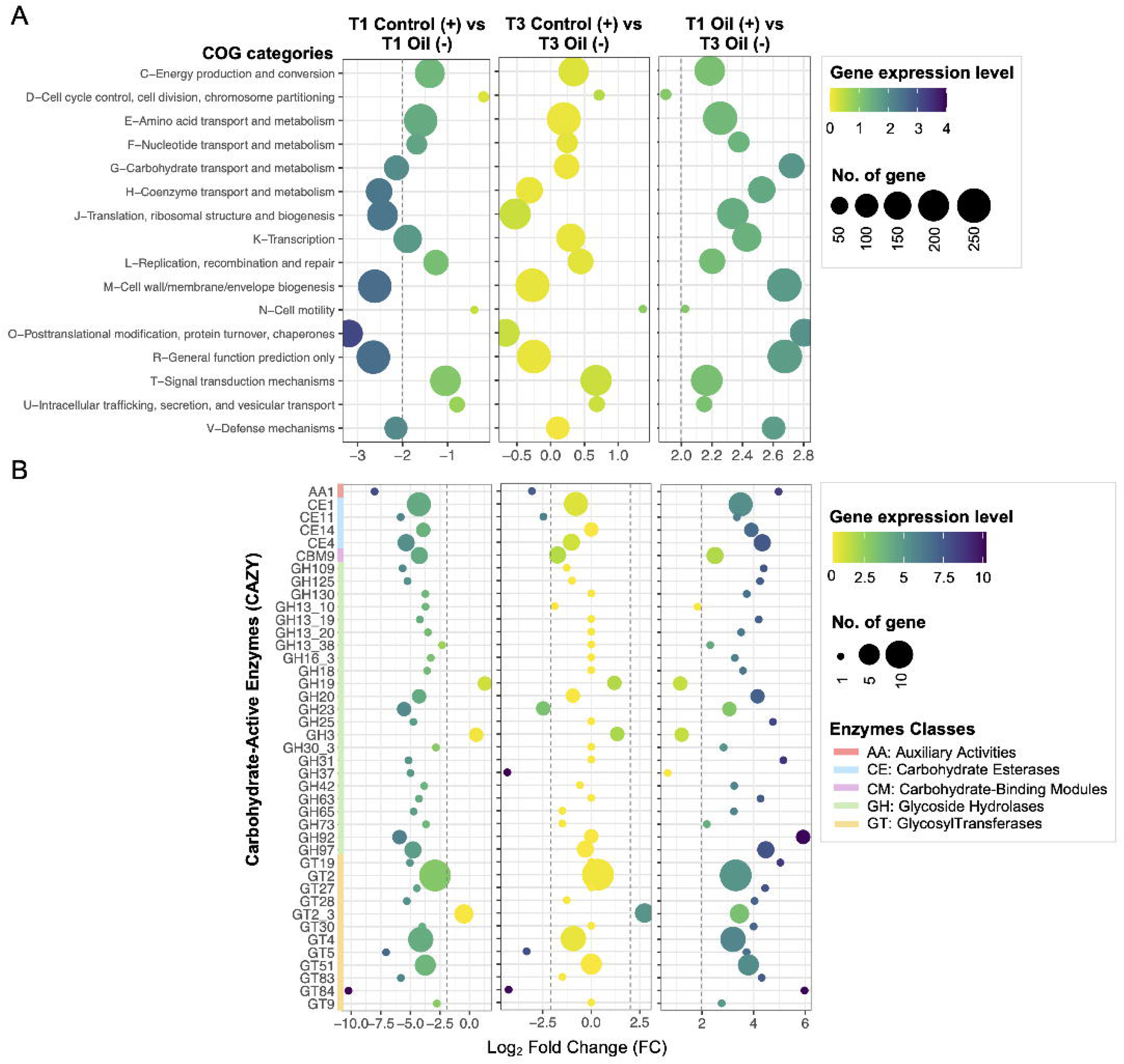
Differential expression patterns of COG categories and CAZymes in response to oil exposure. **(A)** Log2 fold changes of differentially expressed genes across COG functional categories. Dot size indicates gene count (50-250 genes), and color intensity represents expression level (scale 0-4). Three comparisons are shown: T1 Control vs T1 Oil, T3 Control vs T3 Oil, and T1 Oil vs T3 Oil. **(B)** Expression patterns of Carbohydrate-Active enZYmes (CAZy). Families are grouped by enzyme class (AA: Auxiliary Activities; CBM: Carbohydrate-Binding Modules; CE: Carbohydrate Esterases; GH: Glycoside Hydrolases; GT: GlycosylTransferases). Dot size indicates gene count (5-10 genes), and color intensity shows expression level (scale 0-10). Positive log2FC values indicate upregulation, negative values indicate downregulation. Complete gene lists and annotations are available in **Tables S7 and S8.**

To obtain the most comprehensive functional view, we employed Gene Ontology (GO) analysis, which uniquely categorizes genes across three complementary dimensions: biological processes, cellular components, and molecular functions. GO analysis revealed broad cellular responses to oil exposure across the three categories (**Figs. S5-S9, Table S9**). Biological processes showed extensive modulation (log2FC −10 to 5; **Fig. S5**) of amino acid/FA biosynthesis, oxidative stress response, and energy metabolism. Cellular components underwent membrane and peptidoglycan remodeling (log2FC −10 to 8; **Fig. S6**). Molecular functions revealed pronounced changes in oxidoreductases, metal binding activities (iron, manganese), and transport systems (**Fig. S7-S9**), highlighting coordinated adaptation strategies for maintaining homeostasis under hydrocarbon stress.

### Integrated pathway reconstruction reveals cellular adaptation mechanisms

From 5,069 DEGs across all transcriptomes, we selected 181 genes directly involved in hydrocarbon metabolism and cellular adaptation for detailed pathway reconstruction: 108 for cellular response mechanisms (**Fig. 6, Table S10**) and 73 for hydrocarbon degradation pathways (**Fig. 7, Table S11**). Cellular adaptations included oxidative stress defenses (*sodA/N, TrxA/B*) and stress-tolerance mechanisms (*chiA*, a GH18 chitinase), freezing resistance and cold-shock proteins including ice-binding protein (IBP, pf11999) and a GDSL-like lipase/acylhydrolase (pf13472), were upregulated only at T1 (**Fig. 6**). Cell envelope integrity was supported by comprehensive remodeling, including upregulation of lipid A biosynthesis genes (*lpxB/D/H*). Biofilm formation capacity increased through surfactin (*grsT*, cog3208) and cellulose biosynthesis upregulation (*bcsA*, cog1215), accompanied by changes in lipopolysaccharide biosynthesis genes (*lptA/B/D*) and expression shifts of transcription factors *rpoE* (cog1595) and *csgD* (cog2771). Multiple transporter genes also responded, notably the induction of drug efflux systems (*baeS/R, emrA, norM*). Temporal analyses revealed distinct patterns: chemotaxis genes (*cheA/B/R/Y*) and peptidoglycan synthesis genes (*murB/C/F/G)* were upregulated at later stages, while osmoregulation genes (*BCCT*, *mscK/S*,) showed consistent presence under both control and oil conditions. Temperature-sensing systems (*vicR*) and type II secretion (*gspD*) were expressed exclusively under oil exposure.

**Fig. 6.**
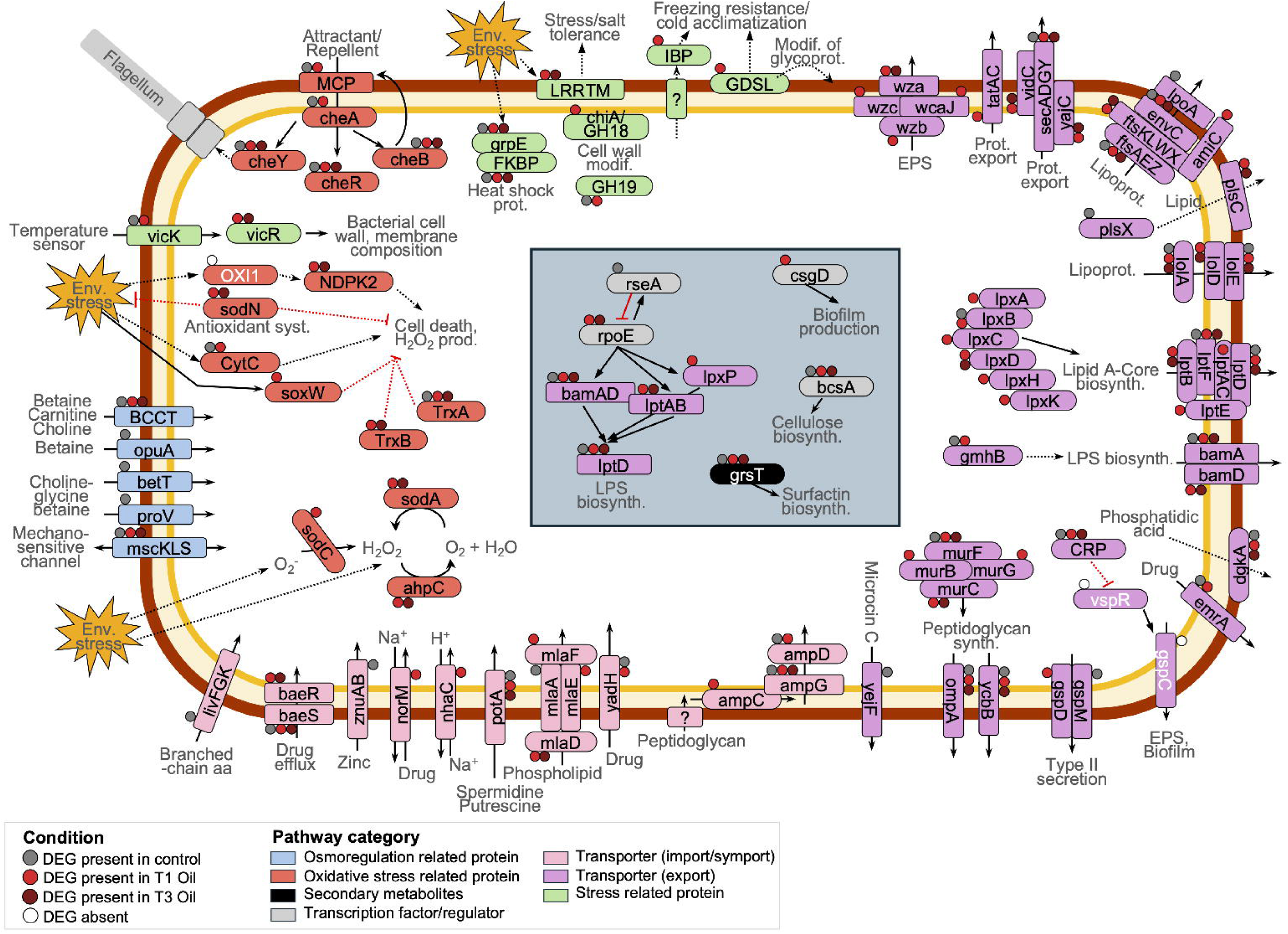
Integrated cellular stress and adaptation responses of *Flavobacterium* sp. R2B_3I to crude oil exposure. Schematic representation of differentially expressed genes (DEGs) mapped onto cellular processes involved in hydrocarbon tolerance. Colored boxes denote pathway categories. A complete list of genes and annotations can be found in **Table S10**.

**Fig. 7.**
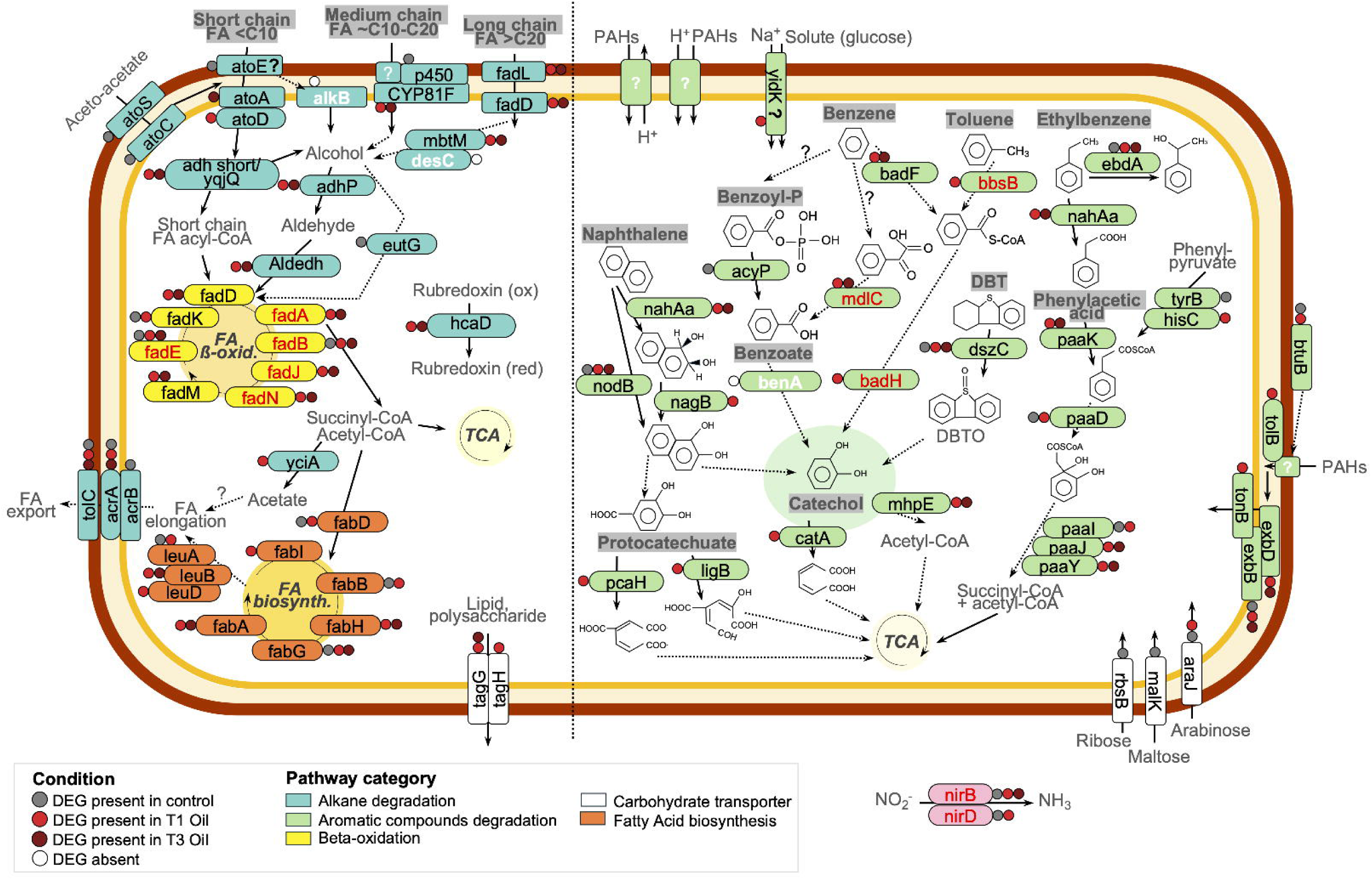
Hydrocarbon degradation pathways and differential gene expression in *Flavobacterium* sp. R2B_3I. Schematic overview of metabolic pathways associated with alkane and aromatic compound degradation during crude oil exposure. Colored boxes denote pathway categories. A complete list of genes and annotations can be found in **Table S11.**

### Non-canonical hydrocarbon degradation pathways in strain R2B_3I

Surprisingly, our detailed analysis of hydrocarbon degradation genes revealed that R2B_3I lacks many canonical alkane degradation systems found in well-characterized hydrocarbon degraders, suggesting it employs alternative metabolic strategies. Several canonical degradation genes, including *alkB* (short-chain alkanes <C10) and *desC* (long-chain alkanes >C20), were absent from the genome (**Fig. S10A, Table S12**). Although the genome encodes a sequence with limited similarity to the *almA* gene of *Alloalcanivorax dieselolei*, phylogenetic analysis positioned it far from the reference, suggesting a functionally distinct enzyme (**Fig. S10B**). Moreover, R2B_3I lacks the *ompW* outer membrane protein typically used for hydrocarbon transport in related species, instead carrying a distantly related peptidoglycan-associated lipoprotein with only minimal similarity (**Fig. S10C**). Phylogenetic and BLAST analyses confirmed absence of conventional alkane monooxygenase pathways present in other *Flavobacterium* species such as *F. xuesshanense*, *F. fryxellicola*, and *F. flevense*. The putative *alkB*-like sequence (hypothetical protein 00363; **Fig. S11**) lacks typical operon organization, missing adjacent rubredoxin genes (*rubA*/*B*), rubredoxin reductase (*rubR*), and regulatory elements (*alkS/R*), and clusters separately from characterized *alkB* genes.

Despite the absence of these canonical systems, the strain expresses alternative degradation machinery, including cytochrome *CYP81F*, multiple *ato* genes (*atoA/E/C/S*), and several *fad* genes (particularly *fadL/D*; **Fig. 7**). Stage-specific expression occurred for *fabI* and *leuD* (FA biosynthesis), *atoD* (*FA* transporter), and *yciA* (terminal step in alkane degradation) at T1 only. Aromatic compound degradation genes also showed temporal specificity: *pcaH* and *ligB* (protocatechuate pathway), *catA* (catechol), *nagB* (naphthalene), *badH* and *bbsB* (toluene), *yidK* (solute transport), *tonB* and *tolB* (PAHs transport), and *hisC* (phenyl-pyruvate) expressed exclusively at T1 oil. In addition, anaerobic degradation potential was indicated by expression of *fadA/B/E/J/N* (for FA degradation), *badH*, *bbsB*, and *mdlC* (for aromatic compounds).

### Novel oxidoreductase repertoire enables hydrocarbon degradation

To identify the alternative degradation mechanisms suggested by the absence of canonical genes, we focused on oxidoreductase enzymes, the primary catalysts of hydrocarbon oxidation, revealing a unique enzymatic repertoire that may represent novel degradation pathways in Arctic *Flavobacterium*. Transcriptomic analysis revealed distinct expression dynamics in response to oil (**Fig. 8, Table S13**). Oil exposure triggered metabolic shifts with 42 oxidoreductases induced exclusively during treatment while 17 control enzymes were repressed (**Fig. 8A**). Of oil-induced enzymes, 23 were expressed throughout exposure while 19 were T1-specific, indicating temporal regulation. Five dioxygenases including 4-hydroxyphenylpyruvate dioxygenase, homogentisate 1,2-dioxygenase, and tryptophan 2,3-dioxygenase maintained expression throughout, while 9 mono/dioxygenases (DOPA dioxygenase, kynurenine 3-monooxygenase, quercetin 2,3-dioxygenase) were T1-specific. Dehydrogenases showed similar patterns with 6 enzymes (alcohol dehydrogenase [NADP+], aldehyde dehydrogenase [NAD+]) expressed continuously and 5 T1-specific. Phylogenetic analysis resolved evolutionary relationships with well-supported clades (bootstrap >50) of alcohol dehydrogenases, CH-CH oxidoreductases, and monooxygenases, highlighting functional conservation underlying R2B_3I’s adaptive hydrocarbon response (**Fig. 8B**).

**Fig. 8.**
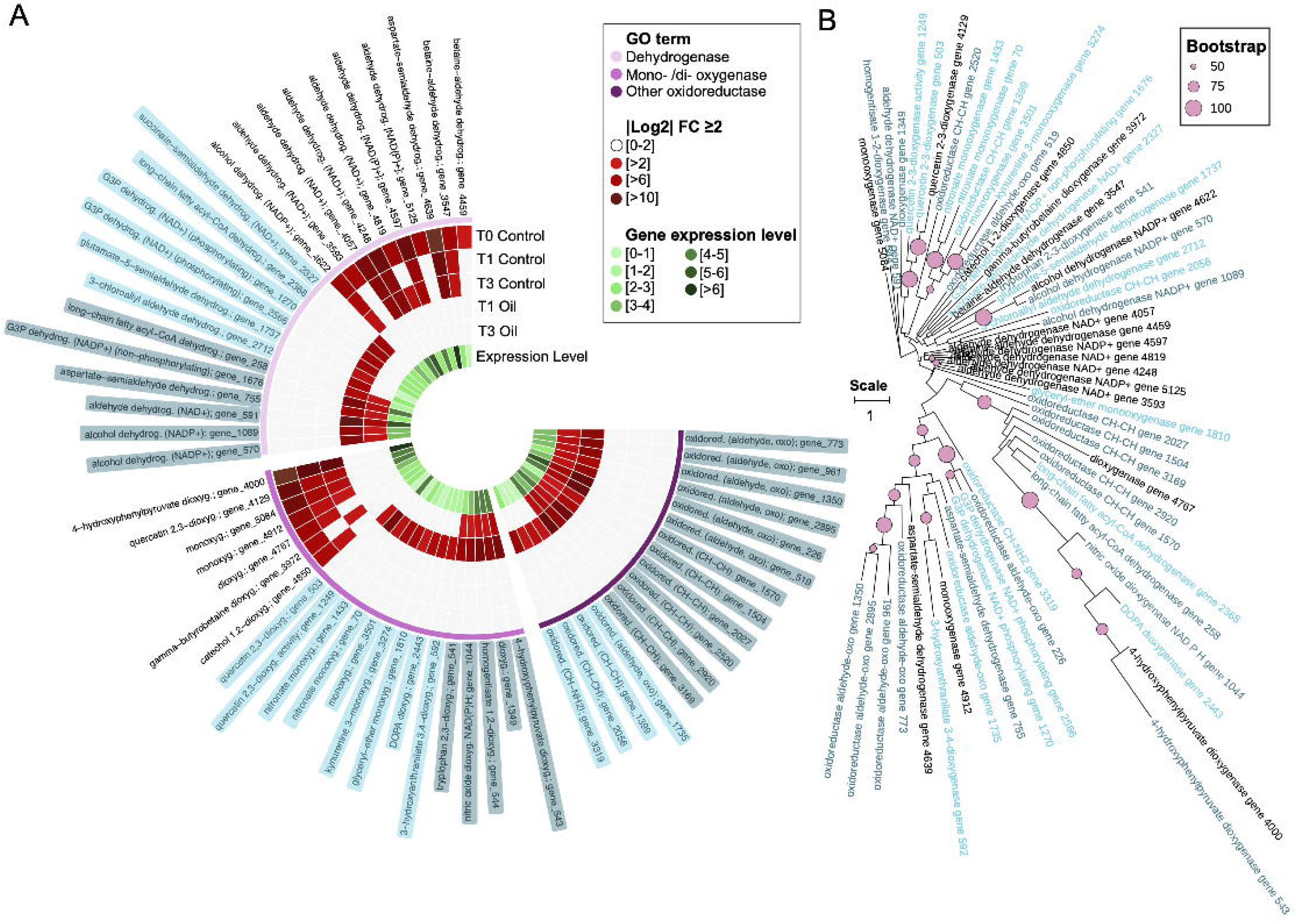
Diversity and expression dynamics of oxidoreductases in *Flavobacterium* sp. R2B_3I during crude oil exposure. **(A)** Circular heatmap of differentially expressed oxidoreductase genes showing functional categories. Inner rings indicate relative expression levels. **(B)** Maximum-likelihood phylogeny of expressed oxidoreductases, with branches colored by enzyme type and nodes marked according to bootstrap support (circle size proportional to support value). The phylogenetic tree was constructed using RAxML from an alignment of 59 sequences, spanning 2940 characters, with bootstrap support calculated over 1,000 repetitions. Only bootstrap values greater than 50 (out of 100) are displayed. A complete list of genes and annotations can be found in **Table S13.**

## DISCUSSION

Our comprehensive transcriptomic analyses of *Flavobacterium* sp. strain R2B_3I reveals a fundamental departure from established hydrocarbon degradation paradigms. Rather than employing canonical *alkB*-type monooxygenases, this Arctic marine isolate utilizes a diverse network of 42 oxidoreductases with distinct temporal dynamics to achieve hydrocarbon metabolism. This discovery, combined with evidence of complex stress management and metabolic flexibility, demonstrates that microbial hydrocarbon degradation encompasses far greater mechanistic diversity than currently recognized. Here, we discuss how these findings reshape our understanding of hydrocarbon degradation evolution, ecology, and biotechnological potential.

### Alternative oxidoreductase networks bypass canonical degradation pathways

The discovery that R2B_3I bypasses *alkB-*dependent pathways while maintaining robust degradation capacity challenges over two decades of consensus that *alkB-*type hydroxylases are indispensable for n-alkane metabolism in marine bacteria [72]. This study presents the first comprehensive transcriptomic analysis of a *Flavobacterium* strain during oil exposure, revealing previously unrecognized degradation strategies that may explain discrepancies between cultivation-based studies and molecular surveys, where *alkB*-negative populations often dominate contaminated sites despite being overlooked by conventional biomarker assays [73–76]. Here, the genome encodes a CYP153 cytochrome P450, which was only expressed under control conditions (**Fig. 7**). Meanwhile, the identified *almA* homologue is phylogenetically distinct from the reference sequences (**Fig. S10**). This suggests that these canonical enzymes are not primarily responsible for oil degradation in this context.

Instead, our data support a metabolic model where alkane catabolism is driven by a non-canonical oxidative network followed by a robust FA processing pathway. In the absence of *alkB* and induced *CYP153*, the initial alkane activation is likely to be performed by the upregulated cytochrome P450 CYP81F (**Fig. 7**). Although P450s are broadly known for their roles in xenobiotic, steroid, and aromatic compound metabolism, many members of this enzyme family exhibit substantial substrate promiscuity. It is therefore plausible that CYP81F functions as a surrogate alkane hydroxylase in this strain. The transcriptomic signature further supports efficient downstream processing of oxidized alkanes: the upregulation of alcohol dehydrogenases (*adhP, adh_short*) and aldehyde dehydrogenases (*Aldedh*; **Fig. 7**) provides the mechanism for converting alkane-derived alcohols into FAs. The induction of long-chain FA transporters (*fadL*), acyl-CoA activation enzymes (*fadD*), and the *ato* gene cluster (*atoA/D*) confirms that these aliphatic substrates are successfully channelled into β-oxidation and subsequently the TCA cycle.

This aliphatic processing operates in parallel with a distinct aromatic degradation strategy. Our model suggests a three-stage process: 1) simultaneous substrate activation via *CYP81F* and diverse monooxygenases, bypassing *alkB* requirements; 2) intermediate processing occurs via aromatic ring-cleaving dioxygenases including homogentisate 1,2-dioxygenase and 4-hydroxyphenylpyruvate dioxygenase that channel metabolites into central pathways; and 3) complete mineralization through the TCA cycle and β-oxidation pathways. This metabolic strategy resembles the biological funneling of diverse aromatic compounds through central pathways in lignin degradation [77], suggesting convergent evolution in complex substrate utilization.

The temporal partitioning of enzyme expression–early monooxygenase induction (T1) for initial oxidative attack followed by sustained dioxygenase activity for long-term processing (T3; **Fig. 8**) reveals temporal metabolic regulation. Early induction is consistent with the role of monooxygenases in oxygen insertion to produce alcohols, epoxides, or diols that activate otherwise inert substrates [78, 79]. By contrast, sustained upregulation of aromatic ring-cleaving dioxygenases indicates long-term processing through central aromatic intermediates including homogentisate, gentisate, and catechol pathways [80]. This redundancy in oxidative capacity, particularly the deployment of enzymes typically associated with aromatic amino acid metabolism (e.g., tryptophan 2,3-dioxygenase, kynurenine 3-monooxygenase), suggests the co-option of existing metabolic machinery for hydrocarbon oxidation rather than *de novo* pathway evolution.

### Genomic plasticity enabling novel solutions

The exceptional genomic plasticity of *Flavobacterium*, characterized by minimal core genome conservation (2.9%; **Fig. 1**) and extensive unique gene content (75.8%), provides the evolutionary framework for alternative degradation strategies. The presence of transposases upregulated at T3 and ongoing L,D-transpeptidase expression (**Fig. S3**) suggests active genomic rearrangement even during prolonged exposure. This architecture contrasts sharply with specialized degraders like *Alcanivorax borkumensis* that maintain conserved degradation clusters [81]. In R2B_3I, the dispersed nature of degradation genes–lacking typical operon organization for *alkB*-like sequences and missing adjacent rubredoxin systems–indicates modular assembly of degradation capabilities rather than acquisition of complete pathways.

The phylogenetic placement of R2B_3I within psychrophilic Clade V alongside *F. gillisiae*, *F. frigidarium*, and *F. psychrophilum,* combined with the absence of canonical *alkB* homologs across the genus (**Figs. S10A, S11**) and only distant similarity to *almA*-like genes (**Fig. S10B**), indicates that cold-adapted *Flavobacterium* species have evolved distinct metabolic strategies shaped by their environmental niche. Such genomic flexibility, observed in other metabolically versatile genera like *Vibrio* and *Pseudomonas* [82, 83], positions *Flavobacterium* as a promising reservoir of novel hydrocarbon degradation mechanisms.

### Integrated stress responses enable hydrocarbon tolerance

R2B_3I exhibits coordinated cellular reprogramming beyond catabolic activation. WGCNA revealed non-random gene organization through modules with strong oil-treatment correlations (**Fig. 2**), confirming biologically coherent regulatory programs. This tight integration of oxidative stress responses with degradative machinery indicates that successful hydrocarbon metabolism requires simultaneous management of toxic intermediates and cellular damage.

The initial stress response phase activates multiple protective systems including superoxide dismutases (*sodA*/*N*; **Fig. 6**) and thioredoxin pathways (*trxA*/*B*) essential for neutralizing reactive intermediates generated during aromatic compound degradation [84, 85]. This protective response extends to structural reinforcement, with lipid A biosynthesis proteins (*lpxB/D/H*) and peptidoglycan synthesis genes (*murB/C/F/G*) showing coordinated upregulation. Furthermore, strong shifts in glycosyltransferases (GT84; **Fig. 5B**) and glycoside hydrolases (GH92), together with persistent expression of laccasses (AA1), and peptidoglycan lyases (GH23), indicate that cell wall modification is critical for withstanding oil stress [86, 87].

COG analysis revealed biphasic regulation of translation and cell wall biogenesis: repression during early stress followed by renewed expression in later phases (**Figs. 5A, S4**). Similar patterns in *Deinococcus radiodurans* following radiation exposure [88], *Rhizobium tropici* under salt stress [89], and *Vibrio cholerae* in acid-tolerance [90], highlight this as a conserved stress-management strategy. Additionally, we observed extensive effects on DNA-related processes, including repair, replication, and methylation. This implies that oil exposure induces genotoxic stress necessitating active genome maintenance, paralleling findings in *Burkholderia xenovorans* LB400 and *Pseudomonas putida*, in which DNA repair systems are upregulated to counter oxidative damage from aromatic catabolism [91, 92].

### Non-canonical degraders reshape Arctic hydrocarbon cycling

Our findings fundamentally alter understanding of marine hydrocarbon fate, particularly in polar regions where *Flavobacterium* represents a dominant bacterial fraction [34–43]. The alternative degradation strategies employed by R2B_3I have profound implications for understanding hydrocarbon fate in cold marine ecosystems. Arctic environments, where seawater temperatures rarely exceed 4 °C and seasonal ice cover creates oxygen gradients, select for metabolically flexible organisms [20–25]. The strain’s potential for anaerobic hydrocarbon metabolism–evidenced by expression of FA β-oxidation genes and candidates like *badH* and *bbsB* under oxygen limitation–suggests it could remain active in stratified water columns and anoxic sediment layers where canonical aerobic degraders fail [49]. Because oxygen limitation is common in contaminated sediments and only a small subset of bacteria (e.g., *Pseudomonas balearica, Desulfoglaeba alkanexedens, Desulfatibacillum alkenivorans*) is confirmed to degrade hydrocarbons anaerobically [93–95], such flexibility would substantially broaden the ecological niche of R2B_3I.

At the community level, R2B_3I’s non-canonical pathways suggest that ecosystem-scale degradation potential has been systematically underestimated. Molecular surveys targeting *alkB* genes capture only a fraction of degradation potential [96, 97], implying that alternative pathway users like R2B_3I constitute a “cryptic majority” that drives substantial hydrocarbon turnover. Given that *Flavobacterium* represents a significant component of Arctic bacterial communities in polar regions, our findings suggest that non-canonical degradation pathways may process a substantial fraction of hydrocarbon inputs in these ecosystems. The widespread distribution of *Flavobacterium* in marine environments, combined with their capacity for horizontal gene transfer, positions them as keystone taxa in hydrocarbon-impacted communities, potentially facilitating community-wide metabolic innovation through mobile genetic elements.

For bioremediation applications, R2B_3I demonstrates that effective cold-water degraders do not have to adhere to traditional metabolic templates. The strain’s integrated stress management systems, including enhanced expression of membrane-stabilizing and cold-adaptation proteins, such as ice-binding proteins and GDSL-like lipases (**Fig. 6**), suggests that cells actively protect envelope integrity during solvent exposure [98, 99], providing insights for optimizing bioremediation conditions. Furthermore, the temporal separation of monooxygenase and dioxygenase expression phases (**Fig. 8**) indicates that biostimulation timing may be critical, with different nutrient requirements during initial attack versus long-term processing phases. Future monitoring must adopt multi-marker approaches or functional assays that capture this full diversity of degradation mechanisms to accurately assess natural attenuation in increasingly accessible Arctic regions.

## CONCLUSION

*Flavobacterium* sp. strain R2B_3I fundamentally challenges hydrocarbon degradation paradigms by demonstrating efficient degradation through alternative oxidoreductases network, entirely bypassing canonical *alkB* pathways. By integrating diverse oxidoreductases with distinct temporal dynamics, oxidative defenses, and membrane remodeling, this strain reveals that hydrocarbon tolerance is a complex, systems-level phenomenon extending far beyond single-gene functions. While specific enzyme-substrate relationships warrant future confirmation through proteomics and defined substrate assays, our current findings have immediate implications for environmental monitoring and bioremediation. Because *Flavobacterium* comprises a substantial fraction of polar marine communities, these non-canonical pathways likely drive significant hydrocarbon processing that is currently invisible to *alkB*-based surveys. This suggests the existence of a “cryptic majority” of degraders and necessitates a revision of monitoring protocols to include alternative markers, particularly as Arctic ecosystems face increasing contamination risks. Ultimately, R2B_3I not only broadens our mechanistic understanding of microbial adaptation but also represents a promising resource for biotechnological solutions in challenging cold-water environments.

## ACKNOWLEDGMENTS

We thank the community of Resolute Bay and in particular Devon Malik for providing logistical support, and his assistance in the field as our guide and bear watcher. We thank Ya-Jou Chen for her help collecting samples in the field. We thank the Canadian Polar Continental Shelf Program for providing logistical support for the field work. This work was funded by Fisheries and Oceans Canada and Natural Resources Canada under the Multi-Partner Research Initiative projects 1 and 2 and by the Fonds de recherche du Québec - Nature et technologies. Arctic logistical support was funded by the Polar Continental Shelf Program from Natural Resources Canada and the Northern Scientific Training Program (NSTP) from Polar Knowledge Canada.

## AUTHOR CONTRIBUTIONS

LGW designed the study. NJF, CG & LGW coordinated the project. AOL conducted field sampling, strain isolation and culture methodologies. NJF and AOL carried out laboratory experiments of liquid cultures. NJF extracted RNA, prepared the cDNA library and analyzed the data. NJF wrote the manuscript. LGW obtained the funding and logistic support. All authors read, revised and approved the manuscript.

## SUPPLEMENTARY MATERIAL

The Supplementary Material for this article can be found online.

