## Supplementary material for "From Northwest Passage shores to molecular pathways: Comparative transcriptomic responses of a novel Arctic marine fuel-degrading *Flavobacterium* species": Fig. S1; Fig. S2; Fig. S3; Fig. S4; Fig. S5; Fig. S6; Fig. S7; Fig. S8; Fig. S9; Fig. S10; Fig. S11

**RUNNING TITLE: Arctic *Flavobacterium* oil degradation transcriptomics**

**Nastasia J. Freyria^1#^, Antoine-Olivier Lirette^1,2^, Charles W. Greer^1,3^ & Lyle G. Whyte^1^**

^1^Department of Natural Resource Sciences, Faculty of Agricultural and Environmental Sciences, McGill University, 21111 Lakeshore Road, Macdonald Stewart Building, Room MS3-053, Ste. Anne-de-Bellevue, Quebec, H9X 3V9, Canada

^2^Graduate School of Agriculture, Hokkaido University, Sapporo 060-8589, Japan

^3^Energy, Mining and Environment Research Centre, National Research Council of Canada, 6100 Royalmount Ave., Montreal, QC, H4P 2R2, Canada

**#CORRESPONDING AUTHORS:**

Dr. Nastasia J. Freyria

21111 Lakeshore Road, McGill University, Ste. Anne-de-Bellevue, Québec, QC, H9X 3V9, Canada

**Table S1 (xlsx).** Overall summary of results of the Illumina transcriptome sequencing.

**Table S2 (xlsx).** Average nucleotide identity values and genome information between selected reference genomes of genus *Flavobacterium* from the pangenome comparison from **Figs. 1 and S7.**

**Table S3 (xlsx).** Petroleum hydrocarbon initial concentration, concentration measured, concentration of degradation and percentage of hydrocarbon removal from **Figs. S1-S2.**

**Table S4 (xlsx).** Statistic one-way ANOVA for total petroleum hydrocarbon analyses from **Figs. S1-S2.**

**Table S5 (xlsx).** List of genes and functional annotation from clusters of top 100 heatmap from **Fig. 2C.**

**Table S6 (xlsx).** List of Kyoto Encyclopedia of genes and Genomes (KEGG) terms from **Fig. 4.**

**Table S7 (xlsx).** List of Clusters of Orthologous Genes (COG) terms from **Figs. 5A and S4.**

**Table S8 (xlsx).** List of genes coding for Carbohydrate-Active enZYmes (CAZY) from **Fig. 5B.**

**Table S9 (xlsx).** List of Gene Ontology (GO) terms, biological process, cellular component and molecular function annotation from **Figs. S5-S9.**

**Table S10 (xlsx).** List of genes and functional annotation found in *Flavobacterium* cell diagram from **Fig. 6.**

**Table S11 (xlsx).** List of genes and functional annotation found in *Flavobacterium* cell diagram from **Fig. 7.**

**Table S12 (xlsx).** Blastp results from comparison of *alkB*, *almA* and *ompW* genes with selected *Flavobacterium* species and other reference species from **Figs**. **S10 and S11**.

**Table S13 (xlsx).** List of genes coding for oxidoreductase enzyme from circular heatmap from **Fig. 8.**

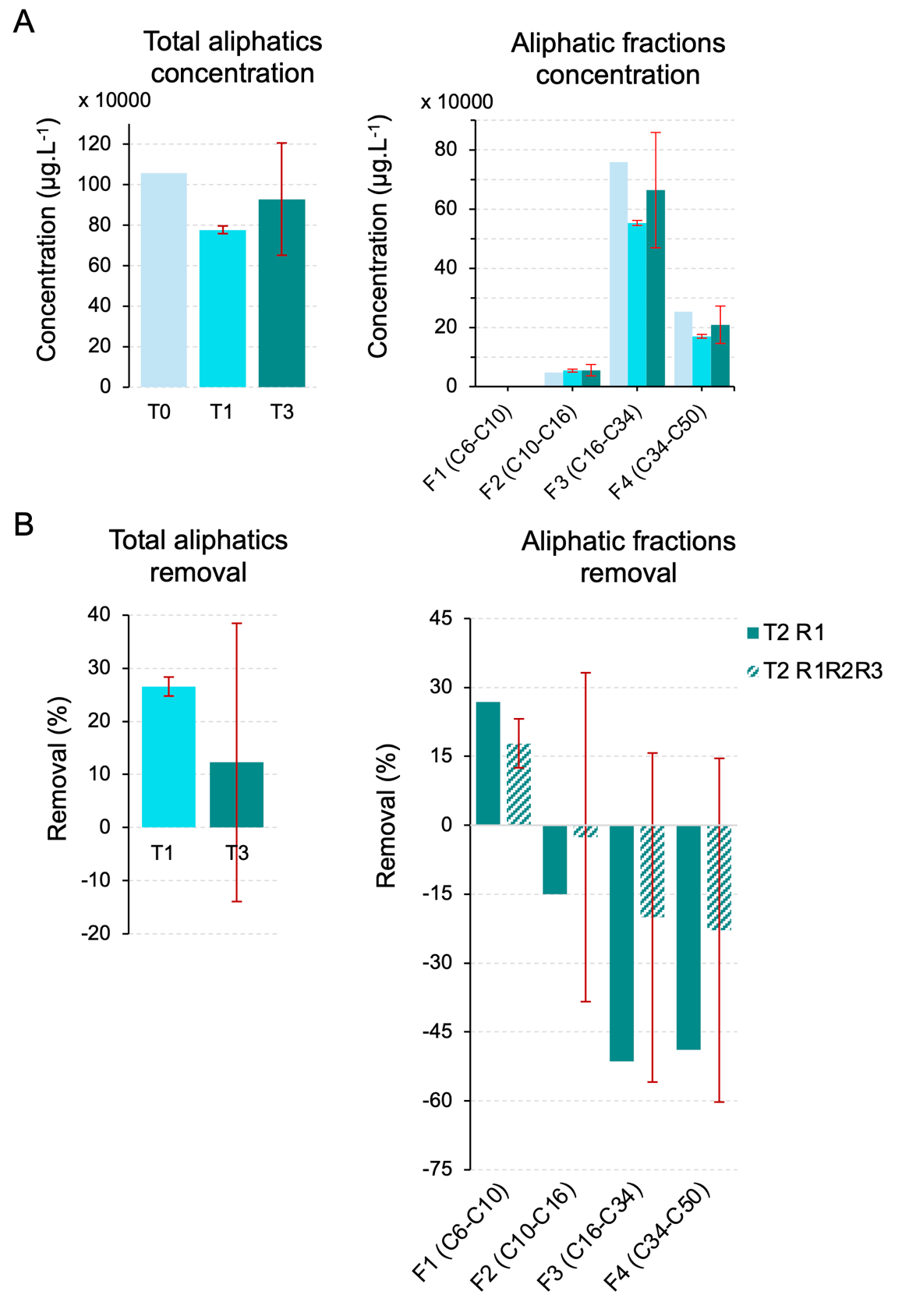

**Fig. S1. Petroleum hydrocarbon analysis during *Flavobacterium* exposure to Ultra-Low Sulfur Fuel Oil (ULSFO). (A)** Alkane fraction concentrations (*µ*g.L^-1^) grouped by carbon chain length, measured at three time points (T0, T1, and T2). F1-F4 represent different carbon chain length ranges. **(B)** Hydrocarbon removal efficiency (%) comparing single replicate (T3 R1, light blue) versus average of all replicates (T3 R1R2R3, dark blue). Red dashed line indicates 80% removal efficiency target. Error bars represent standard deviation (n = 6); negative values indicate compound accumulation or transformation. A complete list of values can be found in **Table S3**.

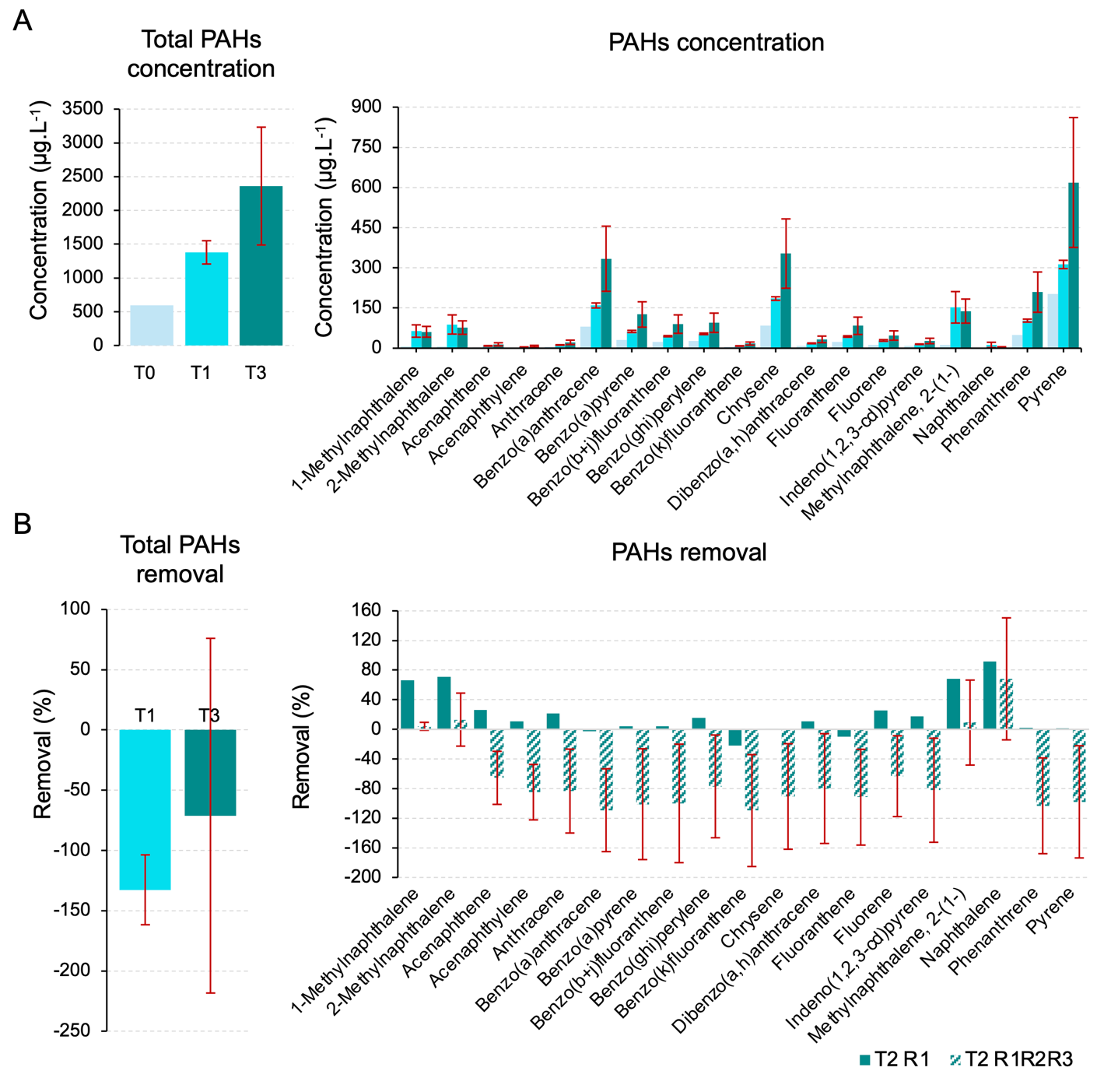

**Fig. S2. Petroleum hydrocarbon analysis during *Flavobacterium* exposure to Ultra-Low Sulfur Fuel Oil (ULSFO). (A)** Aromatic compounds degradation (*µ*g.L^-1^) showing 16 different polycyclic aromatic hydrocarbon (PAH) measured over the same time points. **(B)** Hydrocarbon removal efficiency (%) comparing single replicate (T3 R1, light blue) versus average of all replicates (T3 R1R2R3, dark blue). Red dashed line indicates 80% removal efficiency target. Error bars represent standard deviation (n = 6); negative values indicate compound accumulation or transformation. A complete list of values can be found in **Table S3**.

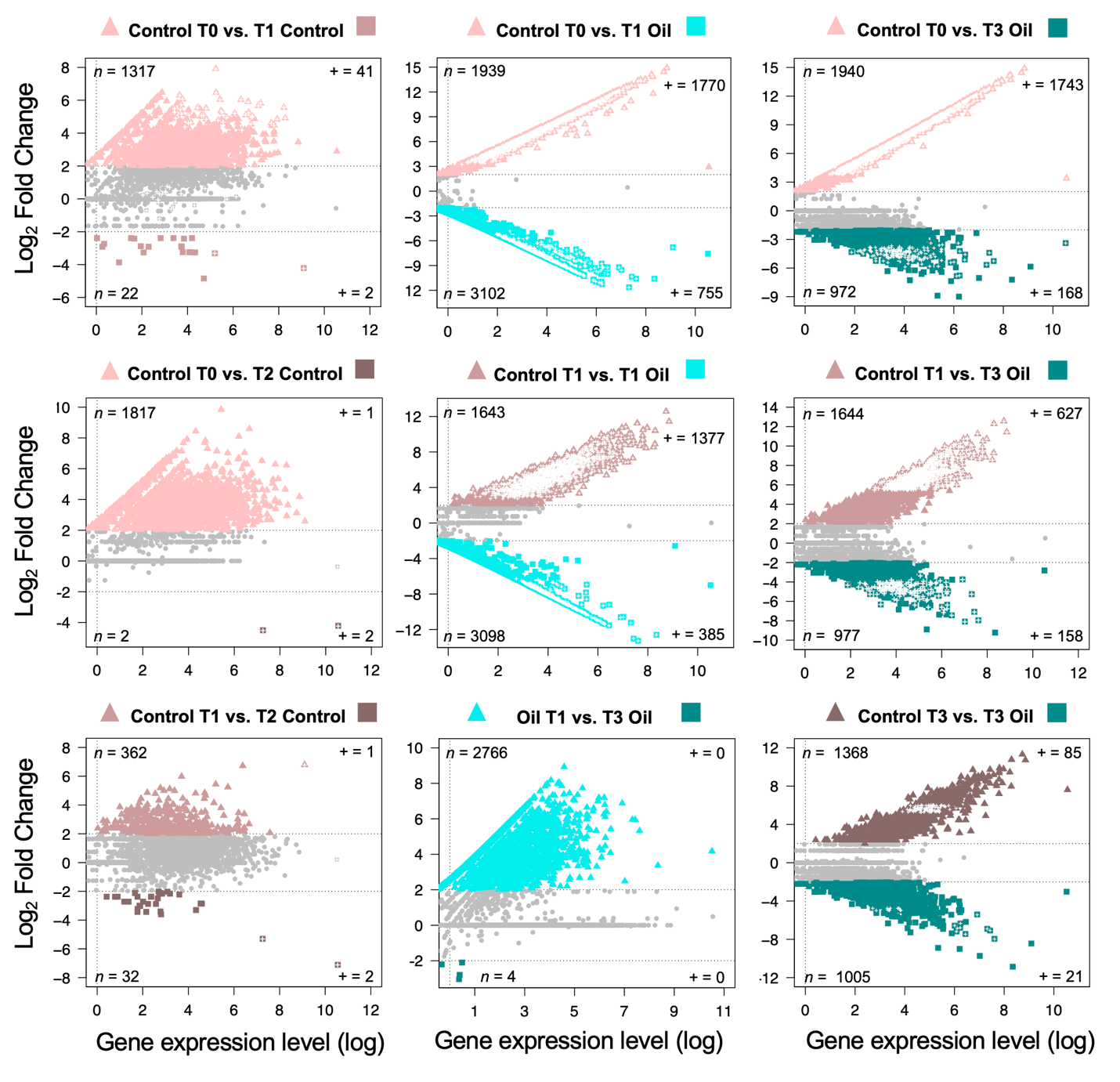

**Fig. S3. Differential expression genes (DEGs) between time of sampling (T0, T1 and T3) with and without oil.** Scatter plots show the comparison between months (T), T1 – 1 month; T3 – 3 months) of cells exposed either with ultra-low sulfur fuel oil (ULSFO) or without oil (control). Each point represents a unigene. Points greater than 2 of log_2_ Fold Change (FC) indicate up-regulated genes and lower than -2 indicate down-regulated genes. Color points indicate differential expressed genes (DEGs). Symbol “+” indicate only filtered DEGs with an Benjamini-Hochberg-adjusted *p*-value <0.05.

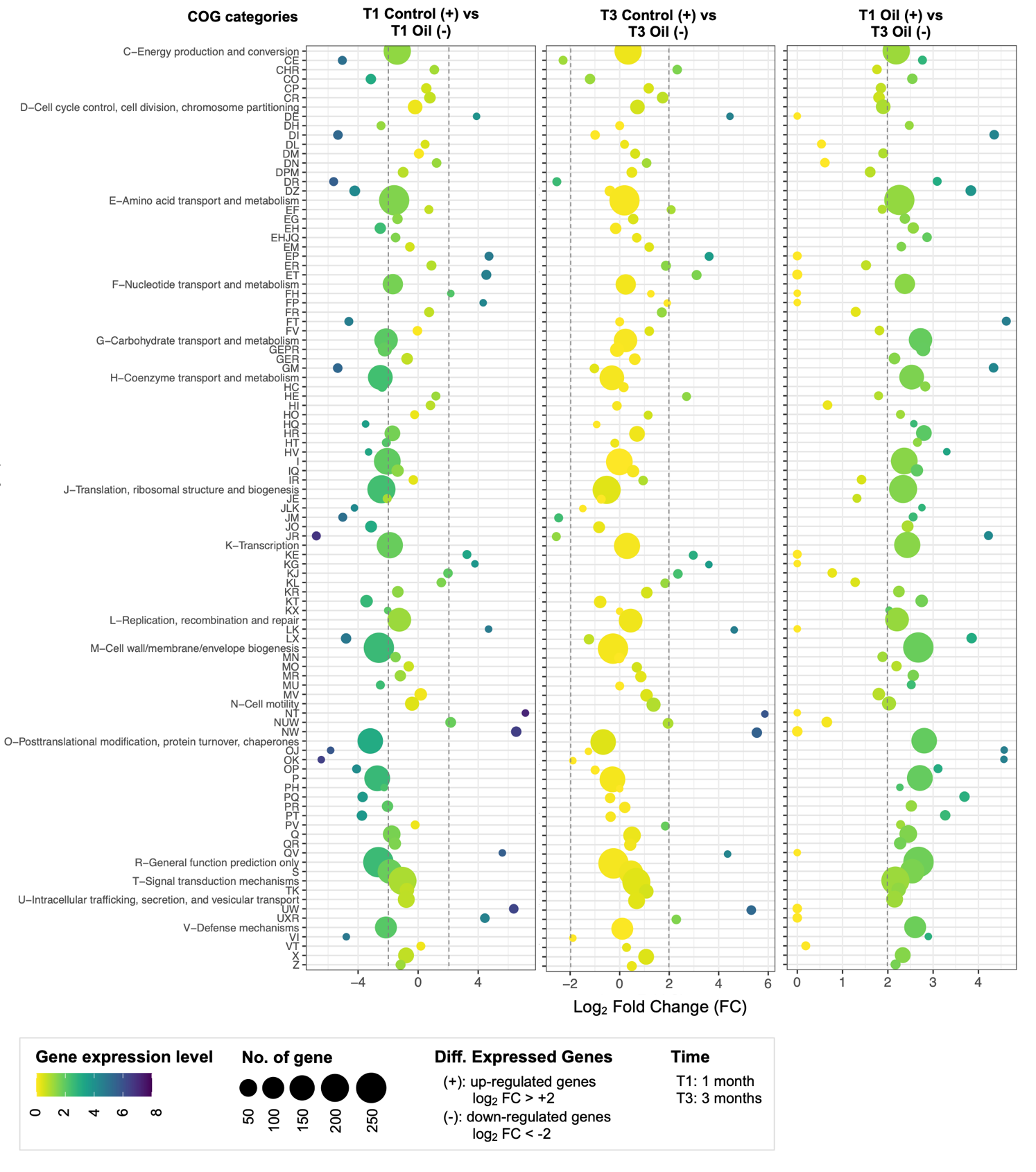

**Fig. S4. Top expression of differentially expressed genes (DEGs) annotated with Clusters of Orthologous Genes (COG) between comparisons of time and with or without the presence of Ultra-Low Sulfur Fuel Oil (ULSFO).** A complete list of genes and annotations can be found in **Table S7**.

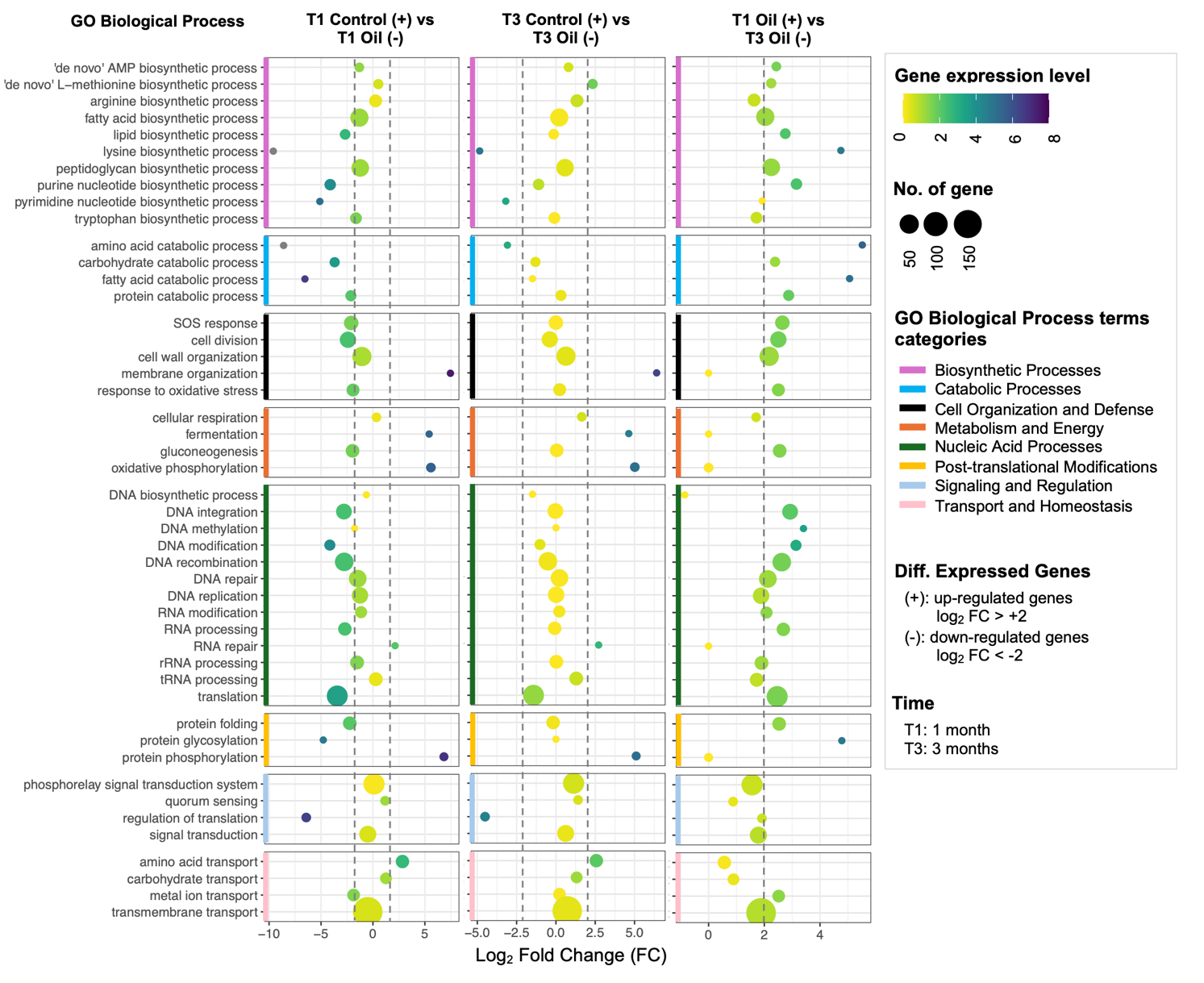

**Fig. S5. Top expression of differentially expressed genes (DEGs) annotated with gene ontology (GO) Biological Process terms between comparisons of time and with or without ULSFO.** A complete list of genes and annotations can be found in **Table S9.**

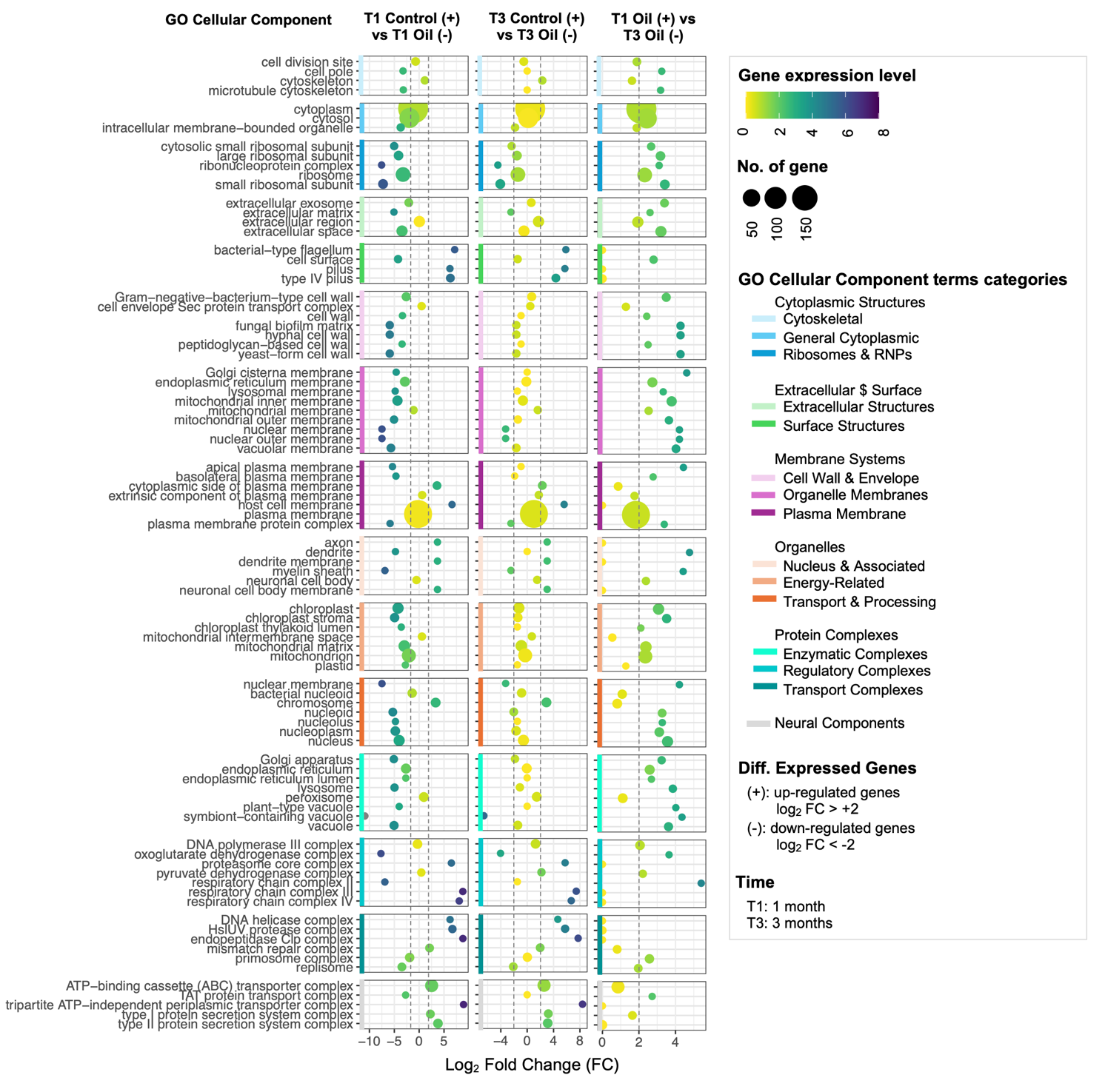

**Fig. S6. Top expression of differentially expressed genes (DEGs) annotated with gene ontology (GO) Cellular Component terms between comparisons of time and with or without ULSFO.** A complete list of genes and annotations can be found in **Table S9.**

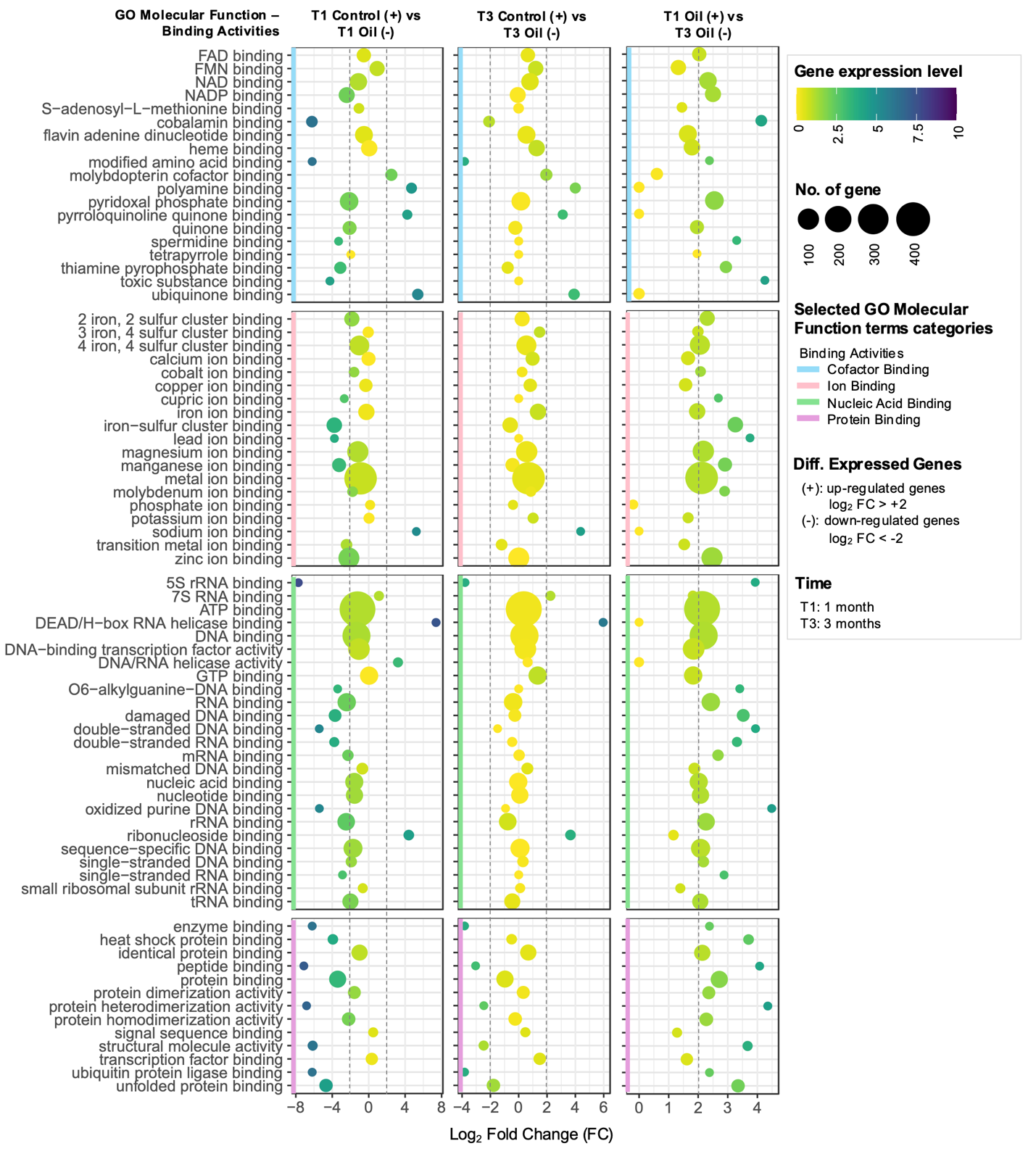

**Fig. S7. Top expression of differentially expressed genes (DEGs) annotated with gene ontology (GO) Molecular Function terms – Binding Activities category between comparisons of time and with or without ULSFO.** A complete list of genes and annotations can be found in **Table S9.**

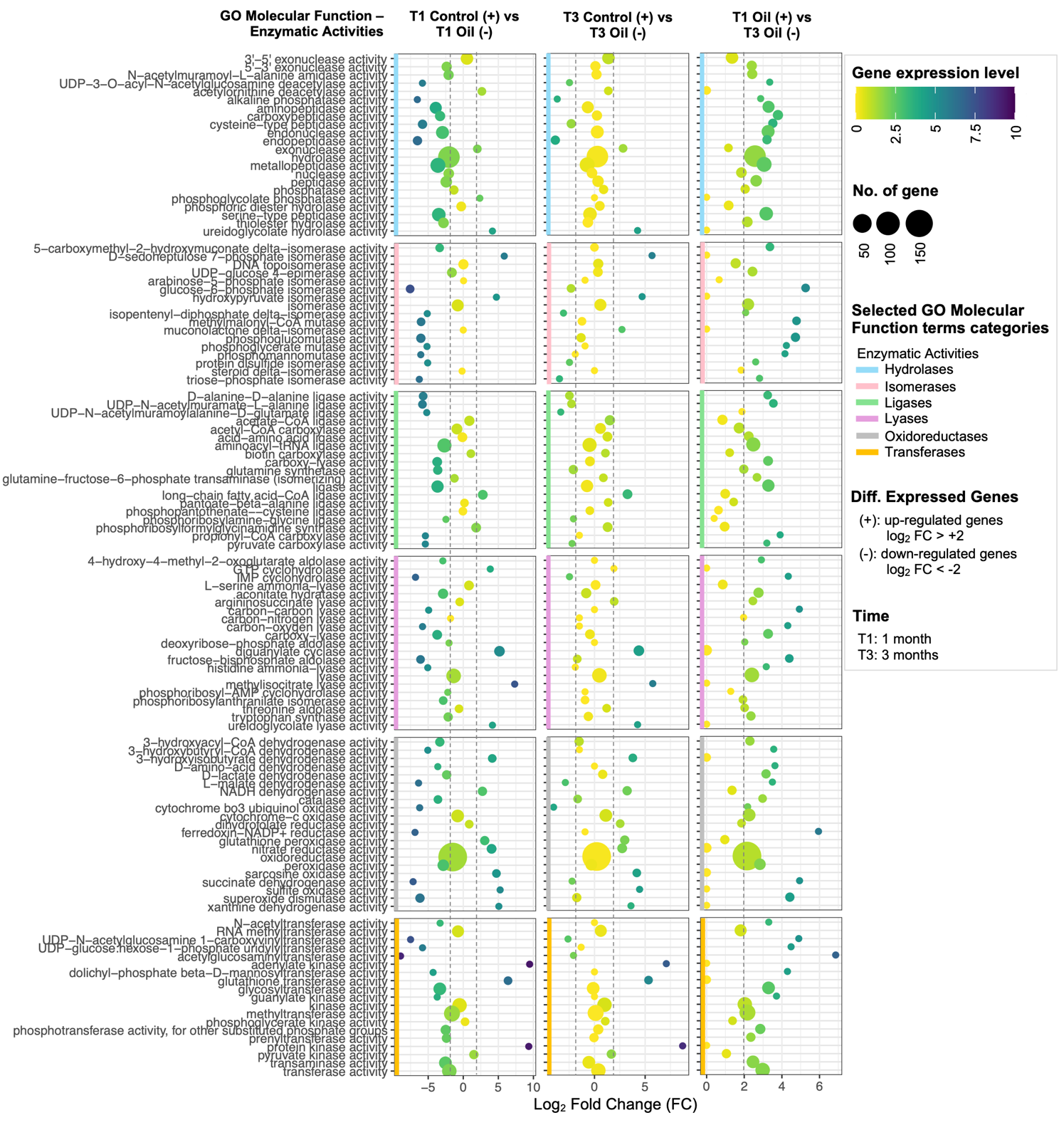
 **Fig. S8. Top expression of differentially expressed genes (DEGs) annotated with gene ontology (GO) Molecular Function terms – Enzymatic Activities category between comparisons of time and with or without ULSFO.** A complete list of genes and annotations can be found in **Table S9.**

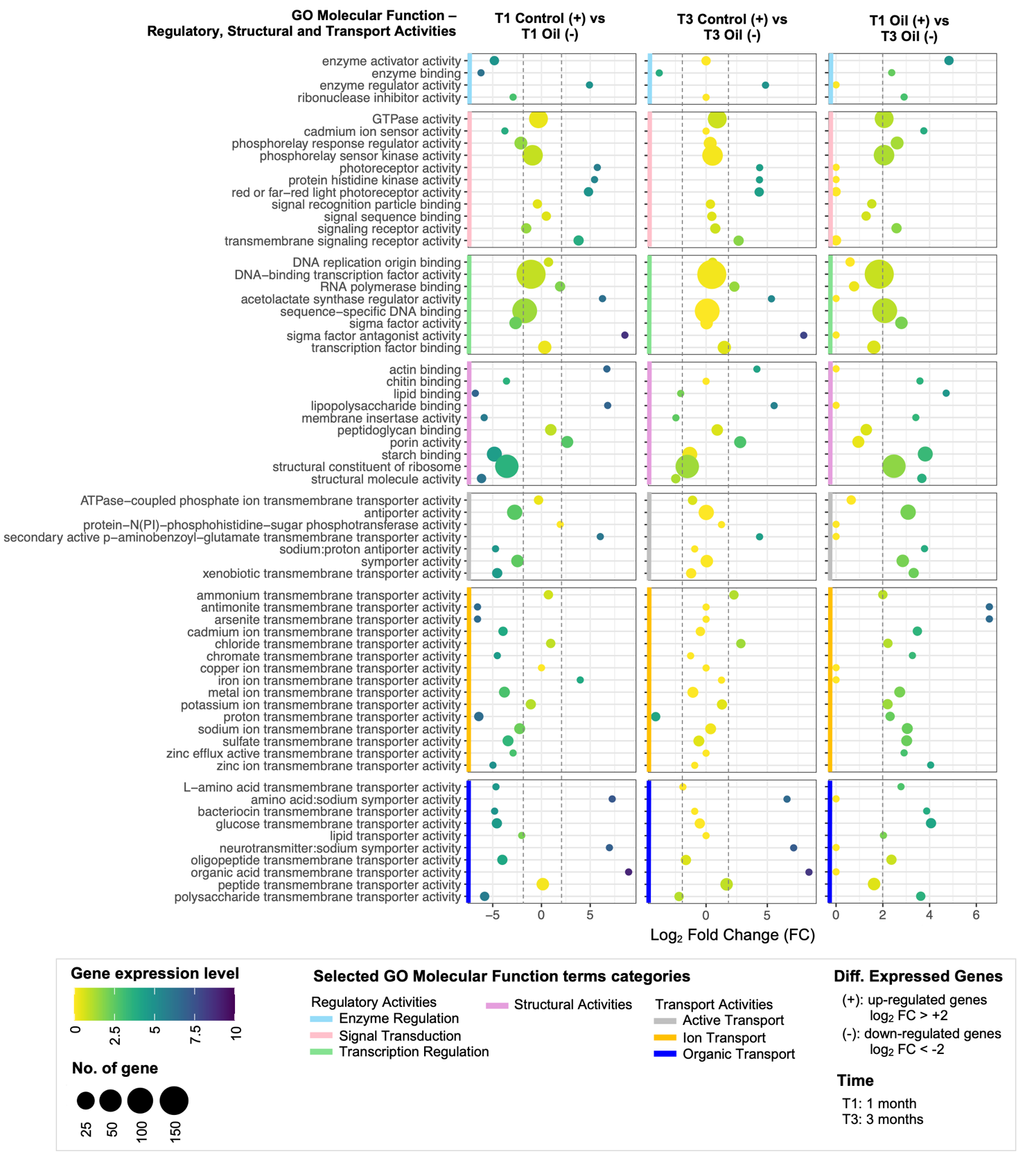

**Fig. S9. Top expression of differentially expressed genes (DEGs) annotated with gene ontology (GO) Molecular Function terms – Regulatory, Structural and Transport Activities categories between comparisons of time and with or without ULSFO.** A complete list of genes and annotations can be found in **Table S9.**

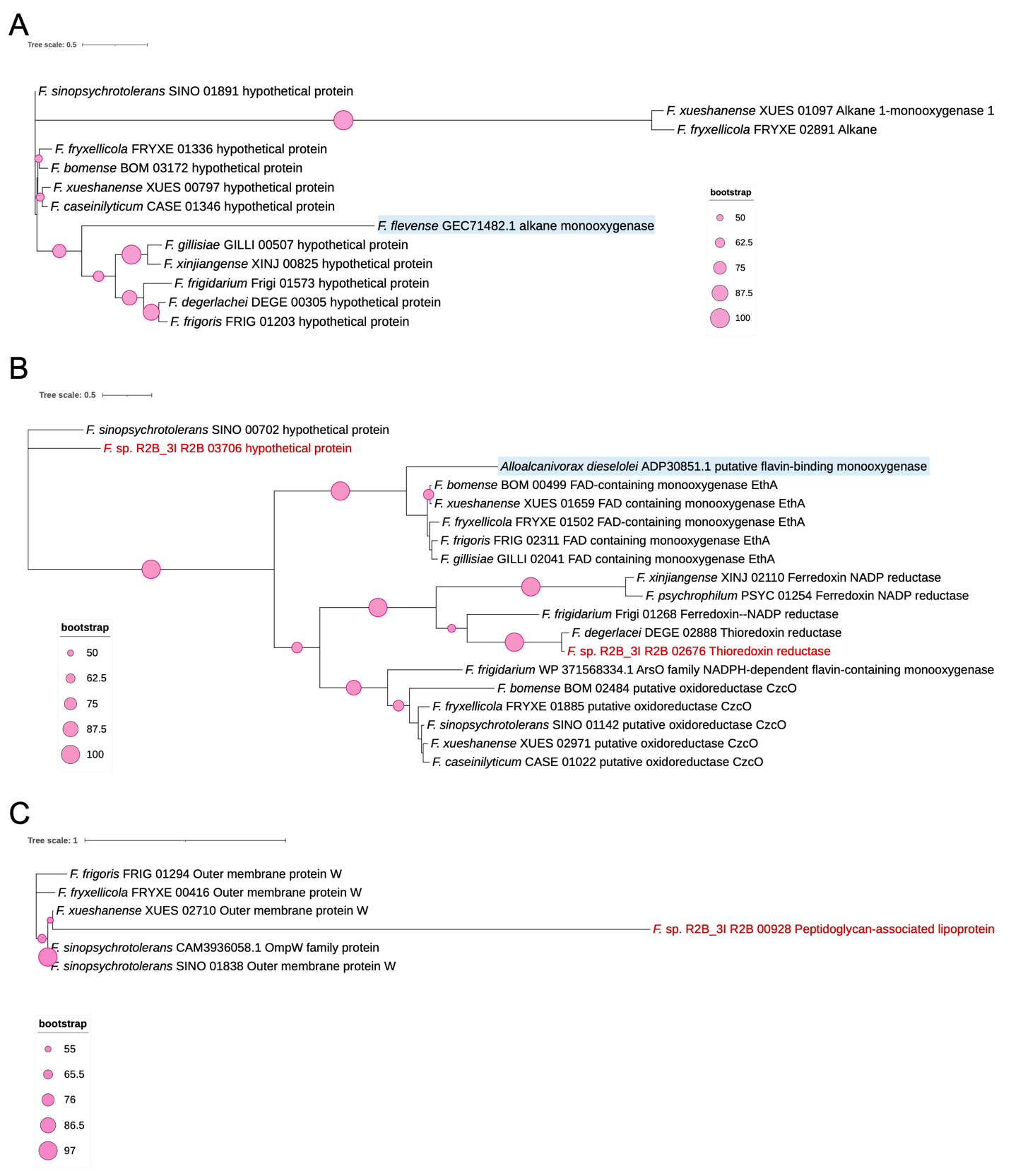

**Fig. S10. A Maximum Likelihood *alkB*, *almA* and *ompW* genes trees generated using RAxML**. (**A**) *alkB* genes phylogenetic tree constructed with 13 sequences or 123 positions of amino acid (aa); (**B**) *almA* genes phylogenetic tree constructed with 19 sequences of 768 positions of aa; (**C**) *ompW* genes phylogenetic tree constructed with 6 sequences of 423 positions of aa. The trees were constructed with bootstrap support calculated over 1,000 repetitions. Only bootstrap values greater than 50 (out of 100) are displayed. Blastp results for *each* gene can be found in **Table S12**.

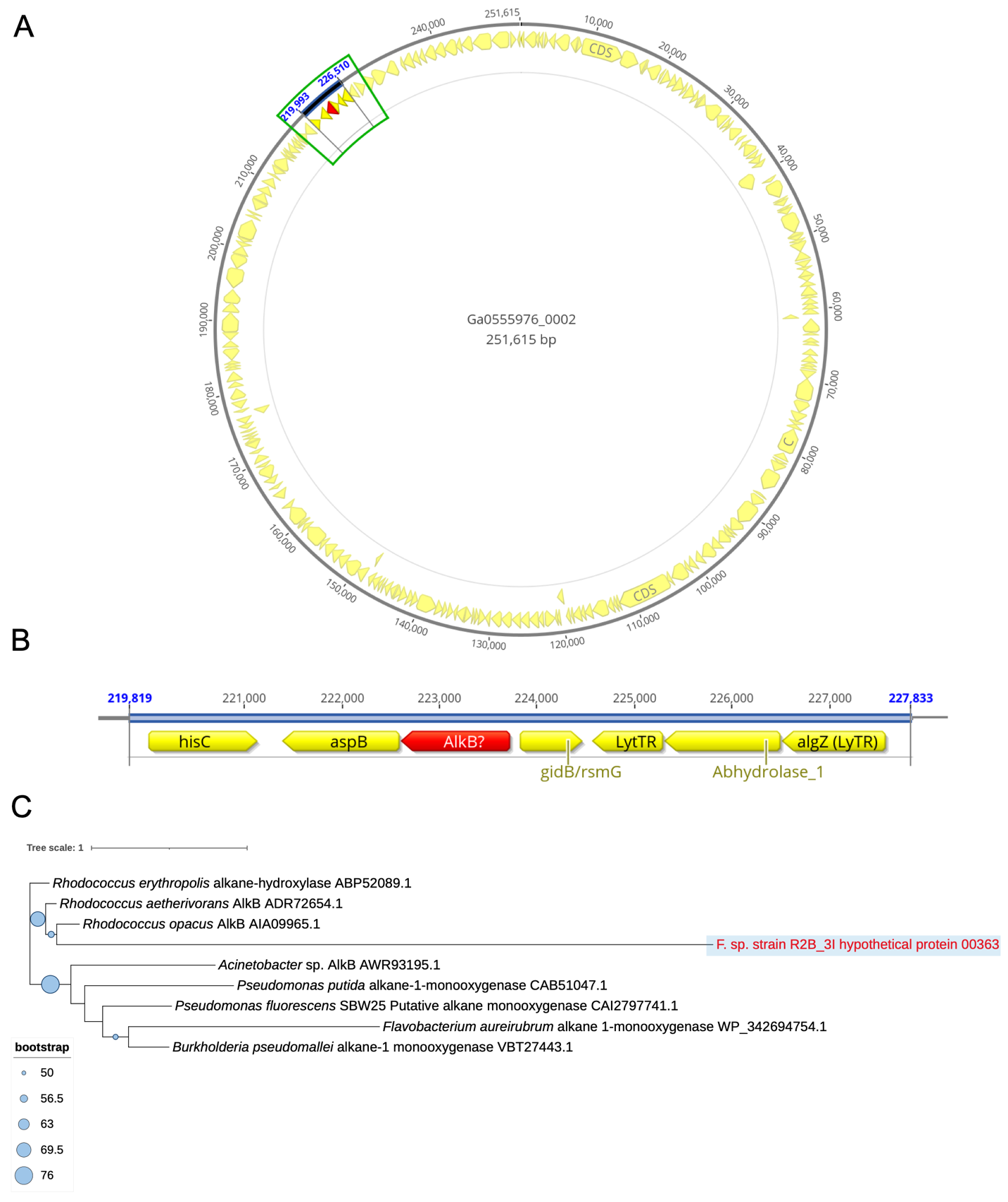

**Fig. S11. Genomic context and phylogenetic analysis of the putative *alkB* sequence in *Flavobacterium* sp. strain R2B_3I.** **(A-B)** Genomic neighborhood of hypothetical protein 00363 showing absence of typical *alkB* operon genes. Unlike functional *alkB* operons that include rubredoxin and rubredoxin reductase genes, this sequence is surrounded by unrelated genes with no hydrocarbon degradation functions. **(C)** Maximum likelihood phylogenetic tree showing the evolutionary relationship between the R2B_3I hypothetical protein 00363 and characterized *alkB* genes from hydrocarbon-degrading bacteria. The R2B_3I sequence (highlighted in red) clusters separately from known functional alkane monooxygenases. Bootstrap values >50 are shown at nodes. Blastp results for each gene can be found in **Table S12**.
